# Inferring the latent network of pairwise mutualistic preferences from observed plant–pollinator interactions

**DOI:** 10.64898/2026.09.17.751682

**Authors:** Leonardo Federici, Eleni Matechou, Iacopo Iacopini

**Affiliations:** Network Science Institute, Northeastern University London, London E1W 1LP, United Kingdom; School of Mathematical Sciences, Queen Mary University of London, United Kingdom; Department of Physics, Northeastern University, Boston, MA 02115, USA

**Keywords:** bipartite networks, hierarchical Bayesian modelling, latent variable models, mutualistic networks, plant–pollinator networks, species abundance, species interaction preferences, urbanisation gradient

## Abstract

Plant–pollinator communities are typically represented as bipartite networks, whose edges are taken directly from field records of visits. These visits, however, are only a proxy for the object of ecological interest: the latent mutualistic preference between two species. While counts are shaped by preference, they also carry confounding factors such as species abundances, sampling effort, and site- or time-specific conditions. We introduce a hierarchical Bayesian framework that treats visit counts as a realisation of a Poisson process and, on the log scale, decomposes the corresponding pairwise rate into a baseline (community-wide activity together with sampling effort), individual species effects representing abundance, and pairwise mutualistic preferences. The model extends to data replicated across sites and time points, and to the inclusion of environmental or experimental covariates. Because the whole system is fitted jointly, we obtain posterior not only for the preferences but for every latent quantity, each carrying ecological signal of its own, with uncertainty propagated through every level of the model, down to any derived network metric. On synthetic data, we show that common practices, such as reading preferences off raw counts or aggregating replicated observations into a single network, confound abundance with preference. In contrast, our framework recovers the underlying preference structure. On empirical datasets, including a seasonal multi-site pollination study where urbanisation level enters as a covariate, the inferred preference network departs markedly from the observed visits, revealing structure hidden in the raw counts: how species vary across sites and time, and which parts of the community respond most to the covariate. When communities are compared along the urbanisation gradient, standard network metrics on the preference layer revise the conclusions drawn from visits alone. The framework offers a principled way to move from networks of observed visits to networks of underlying mutualistic preferences, carrying uncertainty from the data through to the ecological conclusions and accommodating the spatial, temporal, and covariate structure of modern plant-pollinator datasets. Because it acts on the foundational step of network construction, its implications are broad, placing network-based approaches on firmer ground.

## INTRODUCTION

Understanding how species interact within a community drives our ability to manage ecosystems, plan conservation, and predict how habitats respond to perturbation. This is the focus of community ecology, which studies the nature, effects, and ecological context of interspecific interactions. One important class of interspecific interactions is mutualism: a reciprocal relationship in which each species gains a resource or service it could not obtain on its own. Mutualistic systems— communities where such interactions predominate—are both ubiquitous in nature and diverse [8].

Among the most extensively studied such systems are plant-pollinator communities [30], in which plants offer nectar or pollen as a reward, while pollinators carry pollen from flower to flower, enabling the plants to reproduce. These communities are considered instrumental in generating and sustaining the biodiversity we observe on Earth [46], but are threatened by recent environmental changes, and with them the resilience of the ecosystems they support [17, 41].

To make sense of species interactions, ecological communities are often represented as a *network*, with species as *nodes* and *links* (or *edges*) representing a relationship of interest between two species [12]. This approach, rooted in the study of food webs [32], has been naturally extended to plant-pollinator communities. More specifically, a plant-pollinator system forms a *bipartite* network, in which nodes are split into two disjoint sets—here the plant and animal guilds—and links run only across them, wherever a mutualistic relationship occurs (Fig. 1D,F,I). By offering a coarser view, this representation has provided ecological insight into a range of processes, including the effects of species loss [36, 40], restoration interventions [25], and how mutualistic interactions shape stability, coexistence, and resilience to disturbance [4, 13, 45]. Metrics defined on these networks further characterise features not directly observable and make communities comparable, revealing structural patterns that recur across systems [3, 24, 45]. Widely used examples include connectance, degree distribution, nestedness, and indices of how evenly mutualistic relationships are spread across species or across the community [7].

**FIG. 1.**
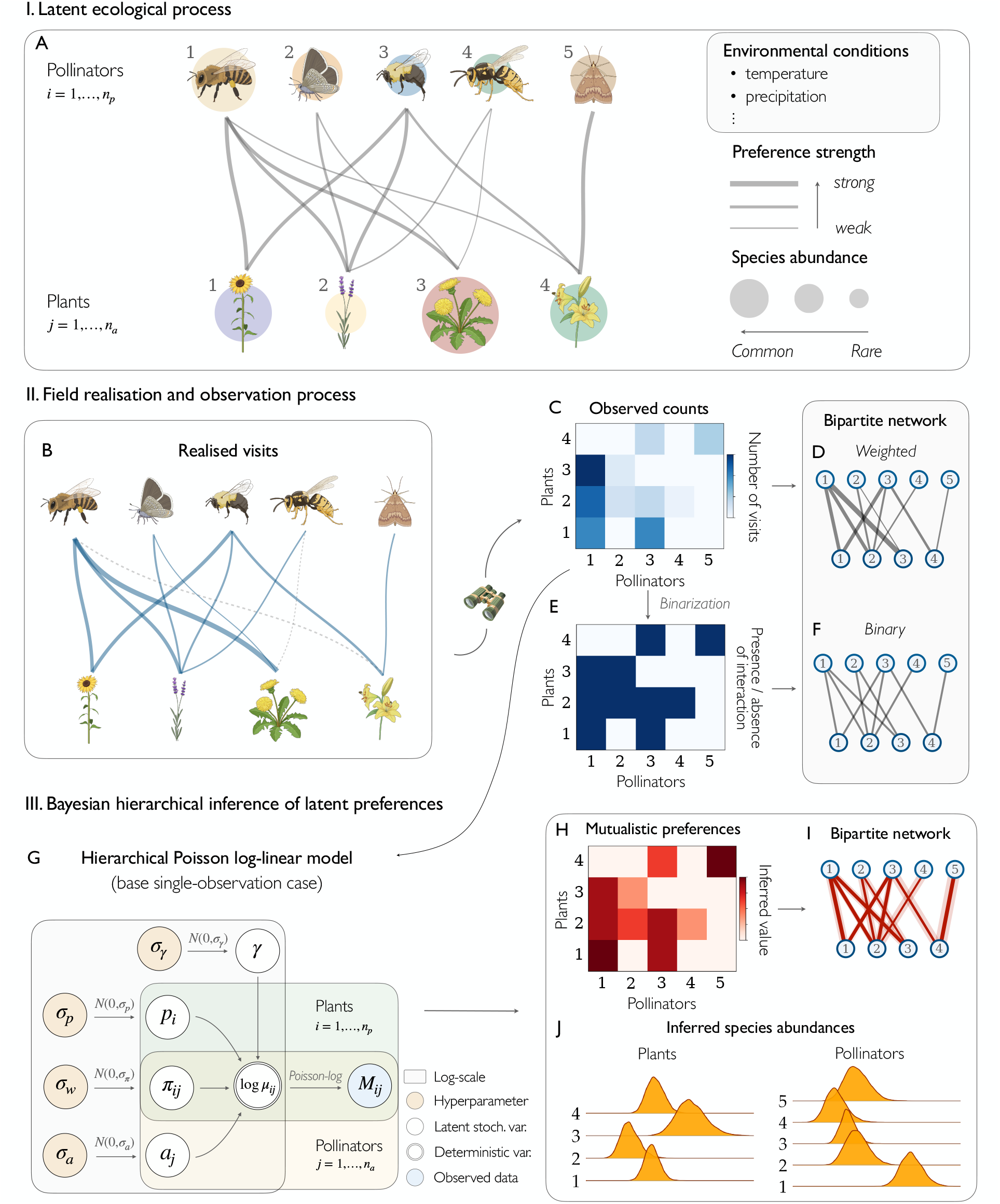
Latent processes behind observed plant-pollinator interactions. (**A**) A community of *n*_*p*_ plants and *n*_*a*_ pollinators, characterised by pairwise mutualistic preferences, individual species abundances and site- and occasion-specific environmental conditions linked to sampling. These factors jointly produce the realised field visits (**B**), which mirror latent preferences but are not identical to them: the mutualism between pollinator 5 and plant 4 appears as relevant as that between pollinator 2 and plant 3, while pair 1–3 shows the strongest apparent relationship driven by abundance rather than preference. Realised visits are sampled into the matrix of observed counts (**C**), with sampling effort introducing further effects. The standard approach transforms this matrix directly into a network of pairs seen interacting (**D**). Counts are often discarded to mere presence/absence (**E**), yielding a binary bipartite network (**F**). Our framework decomposes the log interaction rate into a count baseline *γ* (overall community activity and sampling effort), species effects *p*_*i*_, *a*_*j*_ (abundances), and pairwise preferences *π*_*ij*_ —shown here (**G**) for the single observation case only. Fitting this model to (**C**) provides the inferred preferences (**H**) reconstructing the driving process (**A**). The Bayesian approach returns a full posterior, with shaded regions along edges representing uncertainty (**I**). Posterior distributions are likewise obtained for every other latent variable, including species abundances (**J**).

A crucial aspect of these analyses, however, is how the networks themselves are built. Standard practice constructs networks as mere transcriptions of raw observational data, taking the field record of visits directly as the network. This practice is convenient but problematic for two distinct reasons.

The first is statistical, and stems from the inherent difficulty of ecological data collection [23]. Ecological communities are dynamic, and species interactions vary across time and space [34], so rich, precise, and complete datasets are hard to obtain. Field observations are subject to error and noise, and depend heavily on sampling design and effort, with direct consequences for the networks built on them [20, 37]. Taking the data at face value therefore means leaning on a single, noisy sample of a complex system.

Even with perfect sampling, a second and more conceptual problem concerns how we interpret a link between two species. For plant-pollinator systems the term *interaction* is almost always operationalised in a single way: as a *visit*, an observed event in which an animal is seen on a flower, whose count is read off directly as the weight of an edge. A visit, however, is only a noisy and context-dependent proxy for the *mutualistic preference* between two species, the propensity to associate, shaped by evolutionary and ecological mechanisms. We argue that the object of interest should not be the observed structure of visits but this underlying pattern of preferences. The ecological community has itself repeatedly called for greater methodological caution when analysing observational data, and for a shift toward reconstructing the “true” processes behind observed patterns [6, 11, 47].

Concretely, the visit counts recorded in the field arise from these mutualistic preferences only in combination with species abundances, site- and time-specific environmental conditions, and the community’s overall activity, further shaped by sampling effort [47, 48] (Fig. 1A–C). These counts—that can be seen as the incidence matrix of a weighted bipartite network, or more often collapsed to presence/absence visit records (Fig. 1D–F)—are what researchers usually take as a starting point.

In this paper, we respond to the need for methods that recover the processes underlying observed interactions, and introduce a hierarchical Bayesian framework that reconstructs latent mutualistic preferences from field records of visit counts. Leveraging the flexibility of log-linear models, our method treats the observed counts as realisations of a Poisson process whose rate is factored, on the log scale, into the distinct effects at play, so that preferences can be disentangled from confounding contributions (Fig. 1G). Adopting a Bayesian framework additionally allows us to treat the noisy, error-prone observed network as one stochastic realisation of an underlying process rather than as ground truth. This yields a posterior distribution for each pairwise mutualistic preference (Fig. 1H–I), and more broadly for any quantity derived from the model parameters, from species-level effects (Fig. 1J) and the count baseline to network-level metrics, enabling principled uncertainty quantification for ecological conclusions.

Several recent models share our aim of recovering latent ecological quantities from corresponding visits, but differ in what they take that quantity to be. Staniczenko et al. [44] first argued that an observed weighted network conflates abundance with preference, and proposed extracting the latter, though as a point estimate with no measure of uncertainty. Young et al. [48] work in a generative Bayesian setting, treating visits as noisy realisations of an unobserved network, but their target is binary—whether a preference exists or not—and one preference parameter is shared by every pair; it is recoverable as a special case of the framework we introduce in this paper. The BISoN framework [16] propagates uncertainty in edge weights and corrects for sampling effort, but is deliberately general and does not embed the decomposition into abundance and preference components that we introduce.

We instead infer a separate, continuous preference for every species pair, measured against the visit rate that abundances alone would produce.

Uncertainty propagates from the observed counts to every derived quantity, including the network metrics on which ecological conclusions rest, and the framework accommodates replication across space and time and environmental covariates. Together, these choices bridge two lines of work—uncertainty quantification and pairwise preference inference—within a single Bayesian model grounded in mutualistic plant-pollinator systems.

### Roadmap

We first present the base single-observation model, then its extensions to multiple sites, longitudinal sampling, and covariates. With synthetic scenarios, we test the model’s behaviour and recovery performance under known ground truth. Empirical applications follow the same progression, from a single snapshot to richer multi-observational data, showing the more complete picture our framework provides over raw counts, and the ecological insights that follow. Finally, on a dataset with an urbanisation gradient as covariate, we show which components of the community urbanisation acts on and how conclusions drawn from network metrics on inferred preferences differ from those obtained from raw abundance-confounded visits.

## A HIERARCHICAL BAYESIAN MODEL FOR LATENT MUTUALISTIC PREFERENCES

### From field surveys to interaction datasets

Observational records of plant-pollinator interactions date back to Charles Robertson’s survey of flower visits in Carlinville, Illinois [38], generally regarded as the first such dataset; many more have since been collected and compiled [2]. Two features distinguish them. The first is whether interactions are recorded as binary occurrences or as quantitative counts, typically the number of individuals of each pollinator species seen visiting each plant species. We focus on the latter. The second feature is whether observations extend across *time* and *multiple sites*. This dimension is ecologically critical: seasonal variation, plant phenology, and pollinator life cycles all shape interactions in ways a single snapshot cannot capture [9, 22], and studies increasingly record environmental covariates such as temperature, habitat type, or land use [15, 25]. The model below mirrors this structure directly, from a single-snapshot base case to extensions for sites, time, and covariates.

### Single-observation model

We begin with the simplest form of data usually considered: a matrix of observed counts **M**, with entries *M*_*ij*_ for *i* = 1, …, *n*_p_ and *j* = 1, …, *n*_a_. We follow standard ecological practice for count data and consider each observed interaction *M*_*ij*_ as the realisation of a latent stochastic process:

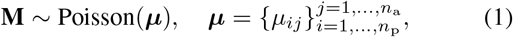

where *µ*_*ij*_ denotes the pair-specific interaction rate, that is, the expected number of visits between plant *i* and pollinator *j*.

The latent process we seek to reconstruct enters through a log-linear decomposition of each interaction rate:

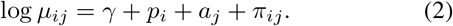

Here *γ*is a global intercept, 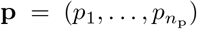 and 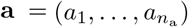 are vectors of species-level effects for plants and pollinators respectively, and *π*_*ij*_ is the pairwise preference between the two species. Exponentiating the linear predictor yields

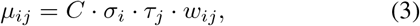

where *C* = exp(*γ*), ***σ*** = exp(**p**), ***τ*** = exp(**a**), and ***w*** = exp(***π***). The constant *C* sets the overall magnitude of counts across all pairs, irrespective of any species-specific effects, absorbing both the community’s intrinsic baseline activity and the sampling effort (i.e., total observation time). The vectors ***σ*** and ***τ*** capture the effect that each species exerts uniformly across all its interactions, reflecting, on the natural scale, species abundances: a highly abundant plant tends to receive more visits from all pollinators, regardless of species-specific preferences. Abundances are rarely recorded in plant-pollinator datasets, so ***σ*** and ***τ***, referred to as “species effects” or “abundances” interchangeably in the paper, serve as proxies for them, absorbing phenological and sampling effects. The preference weight *w*_*ij*_ is read against a neutral value of one (*π*_*ij*_ = 0), where the pair is visited as often as the intercept and species effects predict. Values above one indicate more visits than expected, interpreted as preference; values below it indicate avoidance or incompatibility.

Plant–pollinator datasets vary widely in network size, count magnitude, and sparsity, so we adopt weakly informative priors that accommodate this range without constraining inference [26]. We assign hierarchical priors to the species effects and preferences, parameterised by means and standard deviations throughout:

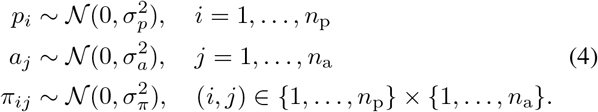

Within each group, the parameters are draws from a common zero-centred distribution whose variance is itself inferred from the data, determining how much species effects and preferences vary across the community. The zero-mean constraint also resolves the additive identifiability issue inherent in Eq. (2): the hierarchical prior anchors each group of effects around zero, with *γ* absorbing the overall mean. The shared scale parameters (*σ*_*p*_, *σ*_*a*_, *σ*_*π*_) further induce *partial pooling* across species and pairs, shrinking rarely-observed species—common in empirical plant-pollinator networks—toward the population mean by borrowing strength from the rest of the community. This provides principled regularisation especially valuable in sparse networks, where individual parameters would be poorly determined. The scale parameters are themselves ecologically informative: *σ*_*p*_ and *σ*_*a*_ quantify heterogeneity in species contributions, *σ*_*π*_ the spread of preferences around neutrality.

Normal priors on the log scale imply log-normal variation on the natural scale, right-skewed and unimodal. For abundances this matches the species abundance distributions observed in ecological communities [29]; for preferences it concentrates mass near the neutral value of one, while the right tail leaves room for the occasional strong affinity—consistent with the view that neutral interactions outnumber strong positive ones in mutualistic systems. Finally, we assign the global intercept a weakly informative prior and place exponential hyperpriors on all scale parameters:

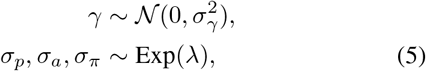

where *σ*_*γ*_ and *λ* are chosen to be weakly informative relative to the expected scale of log-interaction rates.

### Multi-observation model

More commonly, plant-pollinator interactions are observed at multiple sites and sampling occasions. Rather than aggregating these visits—which, as shown below, can lead to misleading patterns—we leverage this granularity within the random effects framework to better infer mutualistic preferences. Data collected at *S* sites and *T* time points (within each site) yield matrices **M**^(*s,t*)^, for *s* = 1, …, *S* and *t* = 1, …, *T*. The observation model then generalises as:

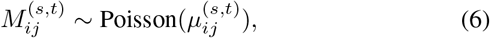

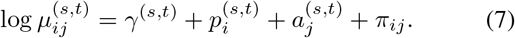

The mutualistic preferences reflect intrinsic compatibility between species pairs, shaped by traits or phenology, and are therefore expected to be approximately invariant across both space and time. Species effects and the intercept, by contrast, can depend on local abundance, phenological timing, habitat features, weather fluctuations, and so should vary across sites and time points. We thus hold *π*_*ij*_ fixed while allowing 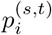, 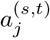, and *γ*^(*s,t*)^ to vary.

The way we let species effects vary rests on a well-established pattern in community ecology: species locally abundant tend to be abundant elsewhere, and vice versa [18], and, by the same logic, they retain a characteristic level of activity across sampling occasions. We therefore decompose each site- and time-specific effect into a species-level *core* and the combined effect of a site- and a time-level *deviation*, instead of treating each 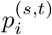as an independent draw from 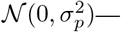 which would discard the shared identity of a species’ effects across observations:

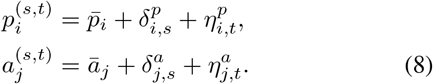

The species-level core and the site-level deviation are assigned the priors

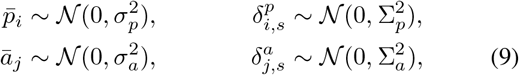

where 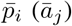 represents the *regional* abundance or activity level of plant (pollinator) species *i* (*j*), its typical contribution to interaction rates across the sites and occasions surveyed, and 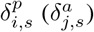 captures its departure from this baseline at a given site due to local conditions. Time-level deviations 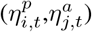 represent species-specific temporal fluctuation, constant across sites: a species generally more active at time *t* is so everywhere. For unordered replicates, such as independent yearly surveys, this can be treated as an exchangeable draw, 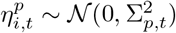. More often, however, time points represent ordered sampling occasions within a season, where exchangeability is ecologically implausible and the process should instead reflect the smooth progression of phenology. In this context, we replace the exchangeable prior with a first-order autoregressive structure:

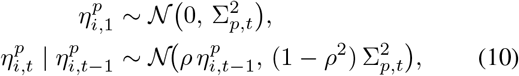

for *t* = 2, …, *T*, where *ρ∈* (*−* 1, 1) is a shared autocorrelation parameter encoding the degree of temporal smoothness, with *ρ* = 0 recovering the exchangeable case. The factor (1 *™ ρ*^2^) keeps the marginal variance constant at 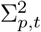 across all occasions, so the process is stationary and 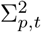 remains directly comparable with the other variance components. This formalises the ecological prior that species activity levels are correlated through the season. We share *ρ* across species and guilds to avoid overparameterisation, though species-specific values could be considered if the data support it. An analogous structure could introduce spatial correlation between sites, mirroring the dependence often encountered among sampled locations.

Finally, for the interaction baseline:

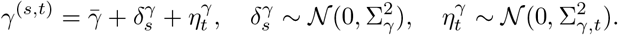

This model formulation naturally includes the special cases of data varying only across space (*T* = 1) or only across time (*S* = 1), where the corresponding deviation simply drops out.

The likelihood depends only on the sums 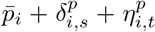 (and analogously for pollinators and the intercept), so the decomposition is not identified by the data alone. As in the single-observation model, identification comes instead from the hierarchical priors, whose distinct distributional assumptions on the core 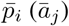 and the site- and time-level deviations partition observed variation into its three scales —more sites *S* and time points *T* providing greater leverage for the separation (see Sec. Model behaviour under simulated data). The corresponding variance components are themselves ecologically informative: *σ*_*∗*_ quantifies *between-species* heterogeneity, while Σ_*∗*_ and Σ_*∗,t*_ quantify *within-species* variability across sites and across time, respectively. Their magnitudes relative to *σ*_*∗*_ indicate how much of the abundance structure is driven by species identity as opposed to local or seasonal conditions.

The same partial-pooling mechanism now operates across sites and occasions, shrinking species observed at few sites or times toward their core, while the random-effects structure keeps the model modular: each additional source of variation enters through its own deviation and variance component, without altering the existing parameters.

#### Adding covariates

The model further accommodates measured spatial or temporal covariates, such as habitat type or temperature, entering the site- or time-level variation in species effects as a component explained by the covariate plus a residual. The same decomposition extends to other model components, like baseline and preferences, as we show later for an urbanisation gradient (see SI Secs. S2, S11).

## MODEL BEHAVIOUR UNDER SIMULATED DATA

### Effect of species-activity heterogeneity and baseline magnitude on preference recovery

Since mutualistic preferences are never directly observable, we first validate the model on synthetic data generated from a known ground truth. We consider three settings: two contrasting single-observation scenarios isolating the effects of species-activity heterogeneity and baseline magnitude; a multi-site example showing the impact of aggregation across times and sites; and a systematic study of recovery across network size, heterogeneity, and baseline visit rate. The first scenario features a heterogeneous pattern of pairwise preferences (Fig. 2A), a homogeneous distribution of species-level effects (Fig. 2B), and a low overall baseline count. Together, these conditions produce an observed count matrix that is itself heterogeneous, with entries of generally low magnitude (Fig. 2C).

**FIG. 2.**
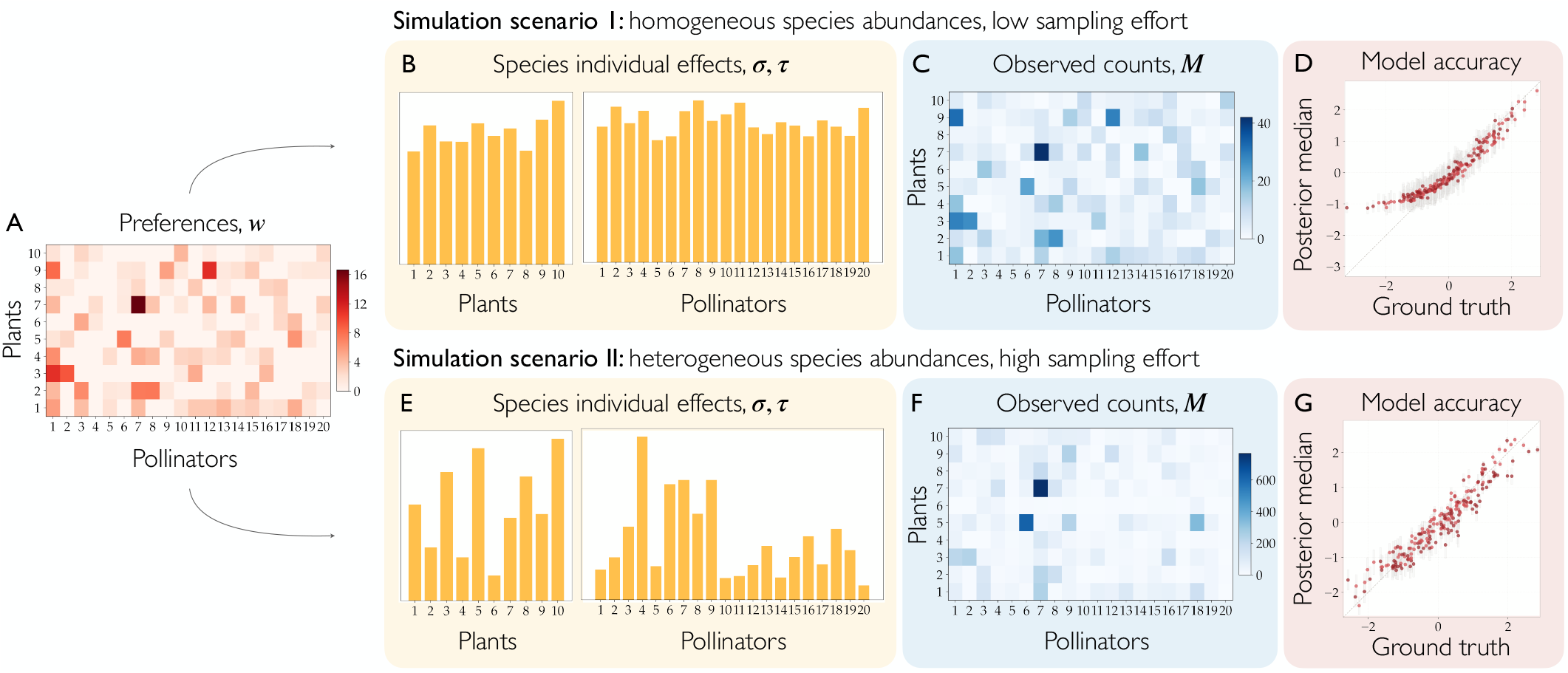
Effect of species-activity heterogeneity and baseline magnitude on preference recovery. Two scenarios share the same heterogeneous pattern of mutualistic preferences (**A**). In Scenario I, homogeneous species effects (**B**) and a low baseline (*γ* = 1) produce the count matrix (**C**); (**D**) compares posterior median log-preferences with the ground truth. In Scenario II, the same preferences are coupled with heterogeneous species effects (**E**) and a high baseline (*γ* = 3), producing the counts in (**F**), with the corresponding recovery in (**G**). The same preferences thus produce different counts under different effects and baselines.

The model recovers most pairwise preferences, yet systematically overestimates those below exp(*−*1) (Fig. 2D). This behaviour is expected: values in this range correspond to neutral or weakly avoidant species pairs, and with limited observational data the model lacks the statistical power to distinguish genuine avoidance from an absence of preference, defaulting to the latter.

The second scenario (Fig. 2, bottom row) is designed as a contrast. Here, the same heterogeneous preference structure is combined with a heterogeneous distribution of species effects (Fig. 2E) and a substantially higher count baseline, reflecting for example a longer or more intensive sampling effort. The resulting observed counts are larger in magnitude and more heterogeneous, while still being close to the patterns seen in published empirical datasets (Fig. 2F). The higher baseline grants inference sufficient power to better resolve neutral and avoidant pairs that proved difficult in the first scenario (Fig. 2G). However, the heterogeneous species effects introduce their own inferential challenge: for preference pairs that are well-identified, the first scenario actually yields sharper and more accurate posterior inference, precisely because the homogeneous effect structure offers a less noisy picture of the latent preferences. The gains of the second scenario are therefore concentrated in its ability to detect the *sign* of the pairwise preferences. This example highlights a strength of the framework: although the same underlying preferences produce markedly different counts as individual effects and sampling effort vary, our model recovers a similar preference pattern, unlike approaches that build the network directly from observed counts or their binarisation (see SI, Sec. S4 for a detailed comparison).

### Aggregating multi-site data confounds preferences with abundance

Aggregating counts across sites discards information and can lead to misleading conclusions. We show this with a synthetic plant-pollinator community sampled at three different sites within the same region. In Fig 3A we report the ground-truth preferences and in Fig. 3B the individual activities of the simulated plants and pollinators, modelled according to our framework as a species core activity plus a site-specific deviation. Coupled with a moderate baseline, these factors produce the per-site observed counts reported in the heatmaps of Fig. 3C-E. The common practice, especially when plant-pollinator networks are used as a basis to simulate the evolution of species abundances over time [13, 39, 40], is to aggregate data across sites, which would result in the matrix of Fig. 3F. Instead, we can fit the multi-observation model to the full data structure. The result is still a single network, but one of inferred latent preferences, obtained by leveraging the complete spatio-temporal resolution of the data. The posterior median of the preferences matrix is reported in Fig. 3G. The model disentangles individual species effects from mutualistic preferences, correctly identifying all 16 pairs of strong mutualism (cf. Fig. 3A), with reduced accuracy for the lower preferences, as expected. Among the top 16 entries of the aggregated counts matrix (Fig. 3F, black outline), only 6 correspond to pairs of (high) preference (red dots). Aggregating over sites thus confounds latent preferences with species abundances, demonstrating the need to disentangle the two. The aggregated observed counts correlate only weakly with the ground-truth preferences (Spearman *ρ* = 0.33, *p <* 0.01) but strongly with the abundance-only null (Fig. 3H, *ρ* = 0.80, *p <* 0.01).

**FIG. 3.**
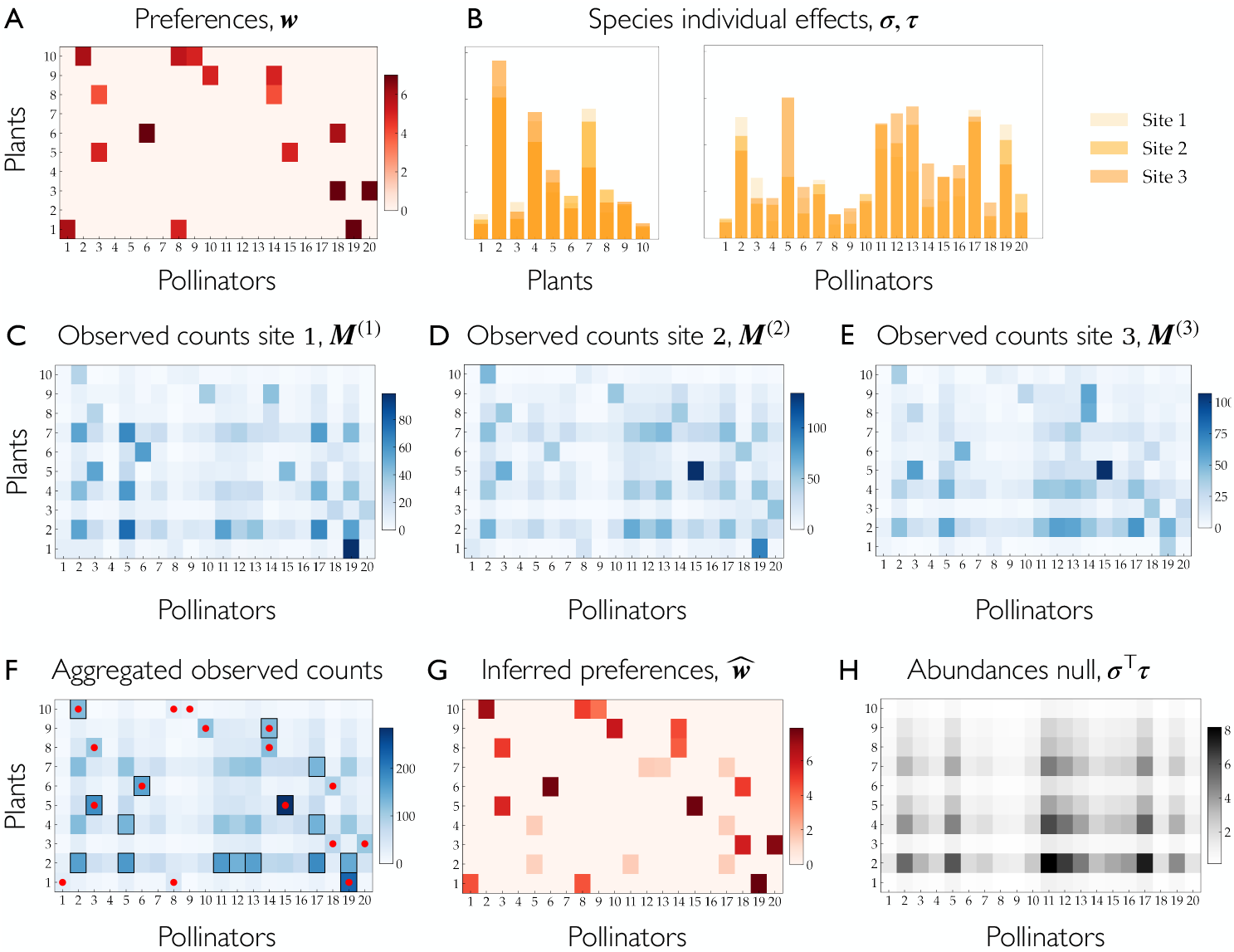
Aggregating multi-observation count data confounds latent preferences with abundances. We simulate a plant-pollinator community with 16 strong mutualistic preferences (**A**), sampled at three sites. Species effects vary around a common baseline across sites (**B**) and, coupled with a moderate sampling effort 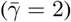, produce the per-site observed counts (**C**–**E**). Aggregating yields the matrix in (**F**), whose 16 largest entries (black outline) include only 6 genuinely high-preference pairs (red dots); the pattern instead tracks the species abundances of the null expectation (**H**). Our model infers the underlying preferences directly (**G**), recovering all 16 strong-mutualism pairs.

### Systematic recovery across the parameter space

Plant–pollinator communities are highly heterogeneous, so to assess how the model performs across a wider range of scenarios, we simulate data sets that vary in network size, species-effect heterogeneity, preference heterogeneity, and count baseline. We fit the model to each and evaluate goodness of fit on the main variables *γ*, **p, a**, and ***π*** through several metrics (details of the metrics and full results in SI, Section S8). Overall, the model recovers the baseline, individual species effects, and pairwise preferences well across conditions, with some loss of accuracy mainly at the extreme values of the parameters. As expected, a higher baseline improves recovery of the log-preference matrix ***π***, consistent with a longer observation time or greater overall community activity providing more information to disentangle pairwise interactions, underscoring the value of well-sampled datasets. We also recover the trade-off seen in the earlier toy example: heterogeneity in species-level effects and the baseline jointly affect the estimation of both the sign and magnitude of a mutualistic relationship (via rank correlation). The toy example could not show, however, that recovery of the baseline itself is degraded by variability in species activity, while it is essentially unaffected by preference heterogeneity. Heterogeneity in pairwise preferences otherwise shows no strong effect, except in the extreme case of very narrow, small preferences (*σ*_*π*_ = 0.1), where the weak signal lowers inference quality. The individual species activity distributions (*σ, τ*) are instead recovered well throughout, degrading only mildly at low baseline combined with high species-effect heterogeneity. Finally, running the sweep at two network sizes yields only a modest improvement with size and no qualitative change.

### Recovery under the multi-site model

How much value do additional sites add? We extend the recovery analysis to the multi-site model with a single network size —justified by the above results (SI, Fig. S10). Performance is good overall, and the site-level deviations in individual species effects (*δ*^*p*^, *δ*^*a*^) are recovered accurately across conditions, largely independent of the number of sites: even three sites suffice, and going to ten changes little. The multi-site setting also lets us ask a more practical question: *given a limited capacity to sample, is it better to observe one site for longer or several sites more briefly?* Comparing the single- and multi-site recovery against the baseline, we find that when per-site counts are low, adding sites markedly improves recovery of the latent pairwise preferences; once the baseline is high, a single, well-sampled site recovers them about as well, and additional sites add little. Notably, this gain from extra sites grows with species-effect heterogeneity; sampling multiple sites helps most precisely when species activity is variable. Given the mathematical exchangeability of sites and temporal replicates, we expect analogous conclusions to apply to repeated, uncorrelated observations over time. We also expect—but do not test here—that correlated time points and mixed time/site sampling would perform comparably or better given the additional information the model could exploit.

Parameter recovery alone does not confirm model validity. Following Bayesian workflow [19], we further stress-test the model with prior predictive checks, calibration, and prior robustness (SI Secs. S5, S6, S7), confirming it behaves as intended when the truth is known.

## EMPIRICAL APPLICATION OF THE SINGLE-OBSERVATION MODEL

We now fit our model to empirical data and compare it against the two standard ways of representing an interaction network—raw counts and their binarisation—to make explicit what our decomposition adds. We start with the single-observation version, fitting it to one of the count matrices reported in the supplementary material of [36]. Its modest size makes it well suited to illustrate both the strength of our approach and the limits of relying on a single snapshot of an ecological community.

Figure 4 contrasts three views of the same community: raw observed counts (Fig. 4A), binarised visits (Fig. 4B), and the quantities our model infers (Fig. 4E-G). The binary matrix discards frequency, treating all interactions as of equal ecological importance. The count matrix retains this information, but conflates two distinct sources: how often a pair interacts because both partners are abundant or active, and how often they interact beyond what their marginal activity predicts.

**FIG. 4.**
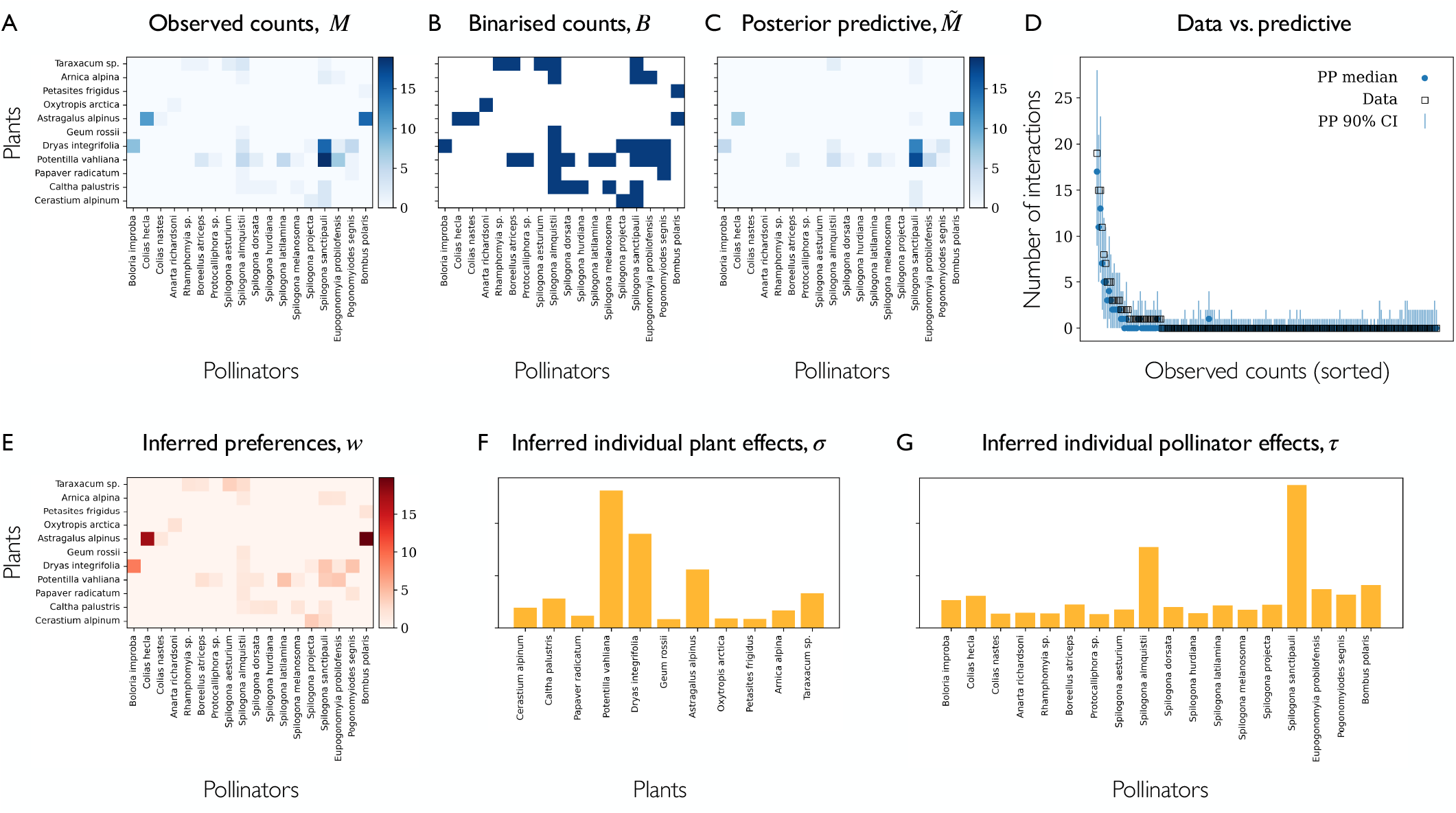
Reconstructing species-level activity and pairwise preferences from a single-observation empirical dataset. We fit the single-observation model to the count matrix labelled as “MOMA” from [36] and contrast three views of the same community. The raw observed counts (**A**) record how often each pollinator species was observed visiting a plant species, while the binarised matrix (**B**) retains only whether a visit ever occurred. Our model decomposes the counts into a pairwise preference matrix (**E**), inferred species individual effects (**F**, plants; **G**, pollinators) and a community baseline (posterior median 2.97, 95% PCI [2.47, 3.44]). Lacking a ground truth, we assess fit through posterior predictive checks: the predictive counts (**C**, posterior-predictive median) recover the observed data well, with some underestimation at the largest values, shown across sorted pairs (**D**; markers denote observed counts, points the posterior-predictive median, and bars the 90% credible interval).

Our model separates the two. The inferred plant and pollinator abundances absorb the first source: a handful of species, notably *Potentilla vahliana* among plants and *Spilogona sanctipauli* among pollinators, carry disproportionately large individual effects, and much of the structure in the raw count matrix (Fig. 4A) aligns with these high-abundance rows and columns rather than with specific pairings. What remains, once these marginal effects are removed, is captured by the preference matrix (Fig. 4E), markedly more concentrated than the count matrix: most pairs sit near the baseline, while a few—the *Astragalus alpinus*–*Bombus polaris* and *Dryas integrifolia*–*Boloria improba* cells among them—stand out as interacting substantially more than abundance alone would explain. Crucially, the cells with the highest observed counts are not, in general, those with the highest inferred preferences. Thus, the raw pattern is driven partly by species-level propensities rather than pairwise affinity, a distinction neither the binary nor the count view can make on its own.

We assess fit through posterior predictive checks (PPC). Propagating the interaction rate sampled at each MCMC iteration and sampling from a Poisson distribution at that rate yields a posterior distribution of counts for every pair. As shown in Fig. 4C-D, the observed counts are recovered well, with some error at the largest values. This first check is *conditional*, meaning each pair is predicted using its own inferred preference, so the reconstruction reflects how well the model fits the given data. The model also lets us examine the inferred variability of the preference weights, and so test whether such a term is needed at all. We obtain a posterior median for *σ*_*π*_ of 2.64, indicating a wide spread of pairwise mutualistic terms across species pairs. Fitting a model without the pair-specific preference and repeating the predictive check confirms this: recovery of the observed counts is markedly poorer, showing that the extension is needed (SI, Fig. S12).

A further strength of the proposed framework is that it infers the rate from which stochastic observed counts were sampled. As a final, more stringent test, we therefore assess the model as a generative process rather than as a per-cell reconstruction. In contrast to the conditional check above, we now integrate out each pair-specific preference, so individual counts are no longer informed by their own observation. What remains driving the predictions is the inferred dispersion 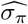, shared across all pairs. For each MCMC iteration we draw a log-preference for every pair from 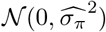, combine it with the corresponding *γ, p* and *a* to form the Poisson log-rate, and simulate new counts. These replicates test whether the inferred dispersion alone 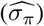 reproduces the variability seen in the data. The mean, the variance, the maximum count and the proportion of empty cells (Table I) all fall close to their observed values, with only a mild tendency to under-disperse. This confirms that the inferred variability of the preference weights is calibrated, and that the pairwise term captures the community’s overdispersion at the right scale. Our analysis, finally, allows us to quantify the uncertainty deriving from relying on a single snapshot, leaving the individual preferences themselves only weakly identified.

**TABLE I.** Summary statistics of the observed interaction counts compared with those generated from the marginal posterior predictive.

| Statistic | Observed | Marginal posterior predictive |
| --- | --- | --- |
| Mean | 0.68 | 0.57 |
| Variance | 5.62 | 4.60 |
| Maximum | 19.00 | 17.67 |
| $P(\text{count} = 0)$ | 0.81 | 0.76 |

## EMPIRICAL APPLICATION OF THE FULL SPATIOTEMPORAL MODEL

### Recovering preferences from multi-site longitudinal data

We demonstrate the multi-observation model on a dataset [25] recording pollinator individuals observed visiting each plant species across 8 sites on Mahé, Seychelles, from September 2012 to April 2013—an ideal test-bed that combines spatial replication with longitudinal sampling. As with the synthetic data, we compare the standard approach of aggregating counts across sites and months against fitting the model to the full spatial and temporal granularity. Figure 5 (for 4 of the 8 sites) shows a substantially different picture. The aggregated counts are dominated by a handful of species in each guild, suggesting—as seen in the synthetic setting—that these simply correspond to the most abundant ones, with observed counts mainly reproducing abundance instead of preference. By contrast, our inferred preferences display a different pattern once individual species effects have been disentangled.

**FIG. 5.**
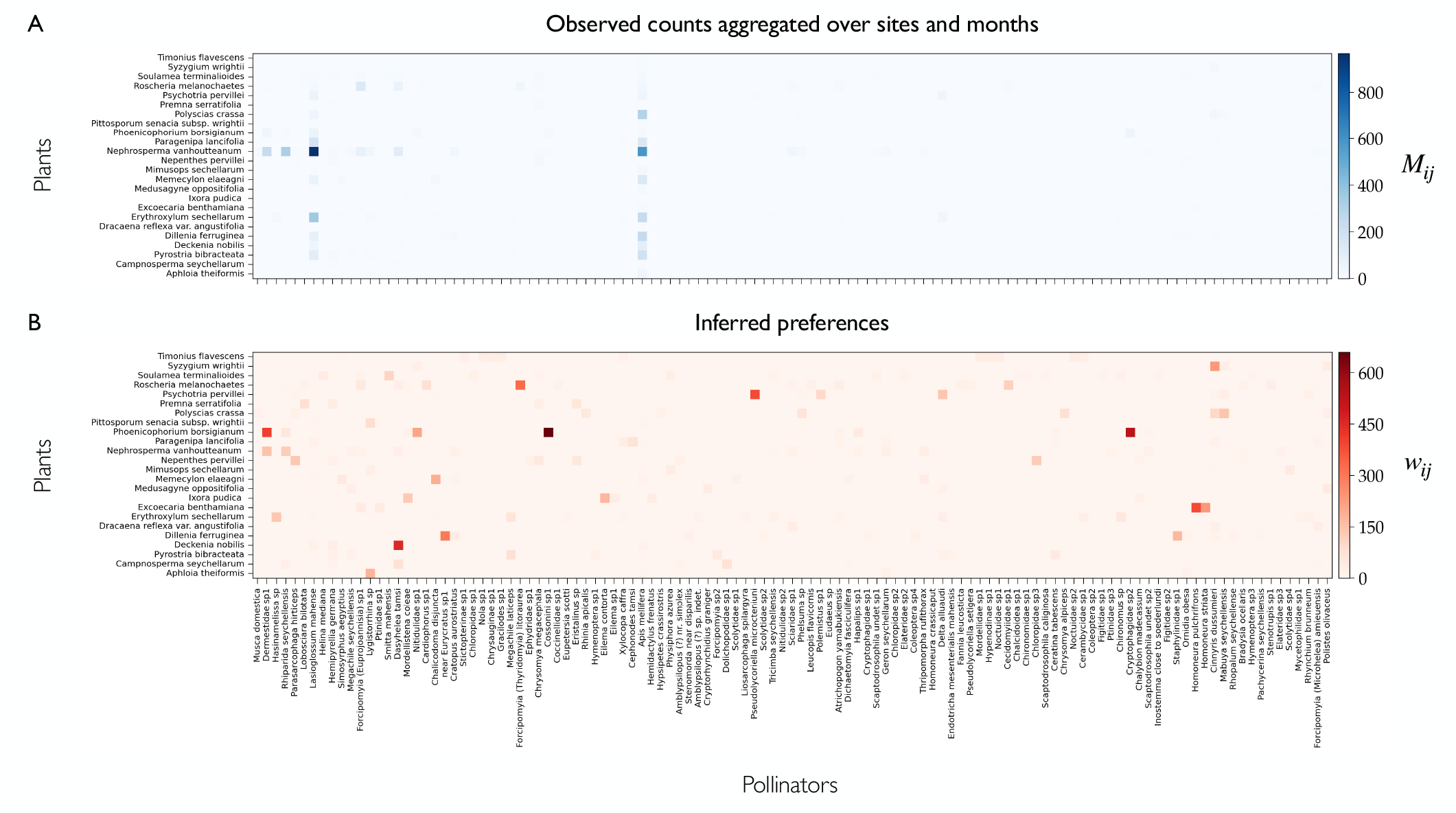
Reconstructing latent mutualistic preferences from multi-site longitudinal data. (**A**) Observed visits of pollinating insects to flowering plants on the island of Mahé (Seychelles), aggregated over 8 sampling months (September 2012 – April 2013) and 4 sites (Bernica, Salazie, Tea Plantation, Trois Frères). Data from [25]. (**B**) Inferred mutualistic preferences (posterior median) fitted by the multi-observation model with correlated time points.

Beyond point estimates, the Bayesian approach returns full posterior distributions for every latent component, each carrying its own ecological signal. The inferred variance parameters (Table II) are particularly informative. Across plant species the core individual activity term is moderate but poorly constrained (*σ*_*p*_*≈* 1.77, 95% PCI [0.05, 3.62]), suggesting plants differ in effective abundance, but this heterogeneity is uncertain, most likely due to sparse plant observations. The pollinator core term, by contrast, is both substantial and tightly inferred (*σ*_*a*_*≈* 1.62, [1.14, 2.03]), indicating markedly uneven individual effects across the guild, a pattern that mirrors the strong abundance skew found in [25]. Site structure separates the two guilds further: among-site heterogeneity in individual effects is large for plants (Σ_*p*_ *≈*2.83) yet modest for pollinators (Σ_*a*_*≈* 0.93). This is coherent with the near-absence of spatial autocorrelation reported for the inselbergs in the original study. Sites differ substantially, but not along a spatial gradient [25]. A plausible explanation is that mobile generalist pollinators bridge differences that patchily flowering plants do not.

**TABLE II.** Posterior summaries from the multi-observation model fitted to data from [25].

| Parameter | Mean | SD | 2.5% | Median | 97.5% |
| --- | --- | --- | --- | --- | --- |
| $\sigma_p$ | 1.773 | 1.016 | 0.053 | 1.873 | 3.621 |
| $\sigma_a$ | 1.616 | 0.228 | 1.144 | 1.622 | 2.030 |
| $\Sigma_p$ | 2.834 | 0.350 | 2.222 | 2.807 | 3.597 |
| $\Sigma_a$ | 0.933 | 0.082 | 0.784 | 0.930 | 1.101 |
| $\Sigma_{p,t}$ | 3.156 | 0.462 | 2.429 | 3.095 | 4.216 |
| $\Sigma_{a,t}$ | 1.496 | 0.161 | 1.239 | 1.478 | 1.872 |
| $\sigma_\pi$ | 2.201 | 0.099 | 2.016 | 2.198 | 2.403 |

Temporal variation is pronounced and, again, strongest for plants (Σ_*p,t*_ *≈*3.16 versus Σ_*a,t*_ *≈*1.50), consistent with flowering phenology and seasonal turnover over the eight-month season. Interestingly, we highlight an asymmetry: for plants the temporal and site terms Σ_*p*_, Σ_*p,t*_ both exceed the crossspecies term *σ*_*p*_ while for pollinators the core *σ*_*a*_ and temporal Σ_*a,t*_ terms are comparable and both exceed the site term Σ_*a*_. Plant activity is thus dominated by phenology far more than by species identity, while for pollinators it seems more governed by species identity and season than by location. Preferences vary strongly across pairs (*σ*_*π*_*≈* 2.20), showing that some interactions are vastly preferred over others beyond the baseline expectation.

### Modelling the urbanisation gradient

Urbanisation affects plant-pollinator interactions [27] and is increasingly recognised as a major global threat to these systems [21], so understanding how it reshapes network structure is essential for planning and conservation. Using the dataset of [15], we ask on which components of the community urbanisation acts, and whether that picture agrees with what raw visit counts alone would suggest.

The study surveyed 12 sites, equally split into low, medium and high urbanisation levels—defined by the percentage of impervious area—in April, May and June. To analyse interaction patterns, the authors aggregated sites within each urbanisation class and treated the 9 resulting level–month networks as distinct communities. Here, instead, we can feed the whole dataset to our model, retaining the crucial information stored in the links between sites, sampling months, and the species observed in each such combination. Focusing on the low and high levels, we introduce urbanisation as a binary spatial covariate *x*_*s*_ ∈ {*L, H*}, agnostically placing a coefficient on every latent layer 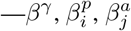 and 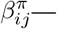 and letting the model reveal which channels show evidence of being affected (for details see SI, Sec. S11).

As before, we examine the community through the model’s posterior over latent variables (Table S6 for details), which is dominated by temporal structure rather than stable species identity. Plant activity fluctuates about twice as much as pollinator activity across visits, consistent with sampling over the flowering season, marking plants as the temporally dynamic side. Preferences, by contrast, are strongly heterogeneous, revealing a marked structure worth tracing along the gradient.

The effect of urbanisation is unevenly distributed across the model: species-specific for plants, near-uniform for pollinators. We show this through the covariate-specific channels (Table III). Community baseline—the only channel free to absorb a net change—shows no credible shift: the slight lean toward fewer interactions per cell at high urbanisation (*β*^*γ*^*∈* [*−*0.63, 0.10]) is not credible, as the interval crosses zero. This agrees with [15], who reported similar total interaction counts across classes, implying that total intensity is roughly conserved among shared species. By contrast, plant species differ substantially and credibly in how they respond to urbanisation 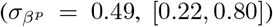, confirming that change is species-specific rather than macroscopic (Fig. S13), while pollinators respond far more homogeneously and weakly 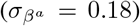. This asymmetry between guilds aligns with the phenological mismatch central to [15], where urbanisation advances flowering phenology but does not shift pollinator flight phenology. Most importantly, we detect a credible and heterogeneous signal of preference rewiring between low and high urbanisation 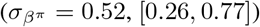, with most pairs barely moving and a meaningful minority rewiring strongly (Fig. S13).

**TABLE III.** Posterior summaries of the covariate-related parameters in the urbanisation multi-observation model, fitted to the dataset of [15].

| Parameter | Mean | SD | 2.5% | 50% | 97.5% |
| --- | --- | --- | --- | --- | --- |
| $\beta_\gamma$ | -0.252 | 0.184 | -0.626 | -0.245 | 0.095 |
| $\sigma_{\beta^p}$ | 0.492 | 0.143 | 0.222 | 0.485 | 0.795 |
| $\sigma_{\beta^a}$ | 0.178 | 0.098 | 0.016 | 0.174 | 0.389 |
| $\sigma_{\beta^\pi}$ | 0.516 | 0.126 | 0.256 | 0.516 | 0.770 |

Beyond individual coefficients, we summarise the effect of urbanisation at the network level using two metrics: weighted connectance and interaction diversity, which captures both interaction richness and the evenness of visits (see SI, Sec. S11D for exact definitions). For each, we contrast three lenses: the posterior distribution computed on our inferred preferences, the aggregated observed count matrix, and the individual site– month networks originally sampled. Weighted connectance shows no substantial difference between classes on any lens (Fig. 6A-B), mirroring [15], who found no difference using binary connectance. Interaction diversity is more informative: the preference posterior separates the classes clearly (Fig. 6C, low 850 vs high 719, *P* (*H < L*) = 0.98), whereas the individual networks overlap almost completely (Fig. 6D, low median 12, high 14, Mann–Whitney test *p* = 0.573). The aggregated counts sit far below both posteriors for both metrics (Fig. 6A, D). Pooling many species into one matrix dominated by a few abundant partners yields a sparse, unevenly filled network that deflates weighted connectance and washes out any interaction-diversity contrast. Only the abundance-controlled preference lens, pooling across sites and months, recovers this difference. Ecologically, this points to a more concentrated, less even preference structure at high urbanisation, compatible with the higher specialisation reported there [15]. The contrast thus depends on the layer examined.

**FIG. 6.**
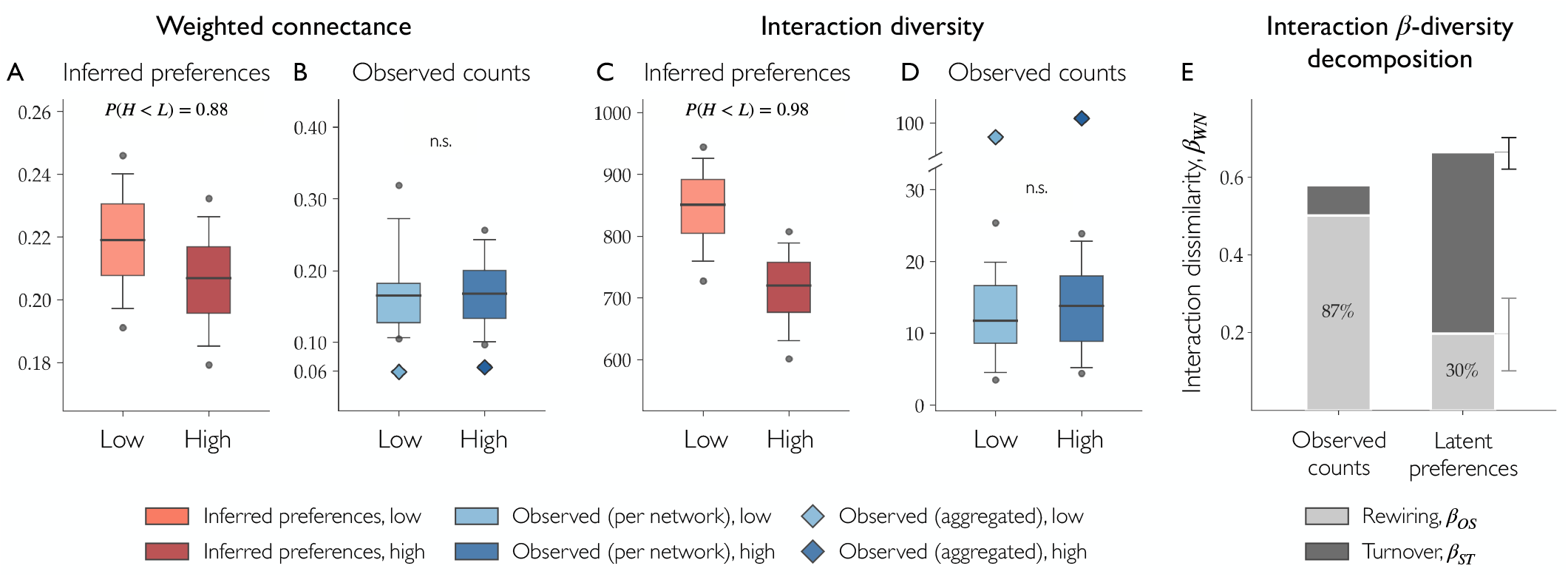
Network metrics contrasts between urbanisation levels depend on the lens. Weighted connectance (**A, B**) and interaction diversity (**C, D**) are compared between low and high urbanisation through the posterior distribution computed on the inferred preferences (**A, C**), the distribution across the individual site–month observed networks (boxes in **B, D**), and the aggregated raw-count network within each level (diamonds in **B, D**). Boxes show the median and interquartile range, whiskers the 10–90th percentiles, and points the 5/95th percentiles. For the posteriors we report *P* (*H < L*), the posterior probability that the high-urbanisation value is lower than the low; for the per-network distributions we report the two-sided Mann–Whitney test (n.s.). **E** Interaction dissimilarity *β*_*WN*_ between urbanisation classes, split into rewiring (*β*_*OS*_, light grey) and turnover (*β*_*ST*_, dark grey), on observed counts and inferred latent preferences. Percentages give the rewiring fraction; error bars, 95% credible intervals.

The interaction-diversity contrast and the rewiring signal in *β*^*π*^ invite a closer comparison with the *β*-diversity analysis of [15]. Using an object close to our interaction diversity (the Hill number of order *q* = 1) and the framework of [31], they concluded that rewiring increased spatial dissimilarity early in the season. Such a dissimilarity, however, conflates three sources of change: species turnover between low and high sites, abundance shifts, and genuine preference rewiring. The authors are explicit that separating the last from the first two was untested in their design. This is precisely the separation our decomposition makes. We therefore adopt the framework of [33], which splits the total interaction dissimilarity *β*_*W N*_ into a rewiring component among species shared by both networks, *β*_*OS*_, and a species-turnover component, *β*_*ST*_. Because our networks are weighted, we measure each term with a quantitative Bray–Curtis index. In addition, we obtain a single season-integrated contrast rather than one per month, as in [15]. Since the seasonal signal is carried by species availability [14], integrating over the season averages out phenological fluctuation while preserving the abundance-free, between-class rewiring the partition targets.

Computed on observed counts, *β*_*OS*_ remains abundance-confounded—two shared species can appear to rewire merely because their activity levels changed. On the abundance-free preferences, that confound disappears and the picture changes substantially (Fig. 6E; credible intervals over posterior draws in SI Table S8): the observed low–high difference looks almost entirely like rewiring (*β*_*OS*_ = 0.50, rewiring fraction 0.87), but on the latent preferences it collapses (*β*_*OS*_ = 0.20) and turnover dominates (*β*_*ST*_ = 0.47, rewiring fraction 0.30). Roughly 60% of the apparent shared-species rewiring is therefore an abundance artefact. The shared species still prefer the same partners but encounter them at different rates, leaving rewiring genuine but minor, consistent with the moderate 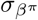 recovered above

A single species pair makes this concrete. [15] highlighted the bee *B. pascuorum* as one of their clearest rewiring cases, visiting *Taraxacum* at low-but not high-urbanisation sites. Yet the urbanisation coefficient for this pair shows no credible effect on their pairwise preference 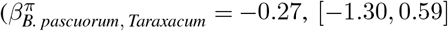, *P* (*<* 0) = 0.72), while *Taraxacum*’s own activity coefficient is credibly negative 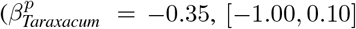, *P* (*<* 0) = 0.90). The apparent partner switch thus reflects the reduced availability of *Taraxacum* under urbanisation, not a shift in mutualistic preference.

## DISCUSSION

We developed a hierarchical Bayesian framework to reconstruct the latent process that drives observed plant-pollinator interactions. Specifically, we decompose the visit count between each pair of species into an overall count baseline, individual species effects, and a pairwise mutualistic preference. These read, respectively, as a sampling effort and community-wide activity level setting the overall count magnitude, as specieslevel abundances (conflated with phenology and other effects), and as the underlying mutualistic affinity between pairs, independent of site, time, and sampling-specific effects. This shifts analysis from the network built directly from observed data—or, more commonly, its binarised version—to a weighted network whose links reflect the latent mutualistic preference. In doing so, we respond to recent calls for a sounder methodology when building networks from plant-pollinator observations, and for a shift towards reconstructing the latent processes that generate the visits we record [6, 11, 47]. Crucially, the Bayesian approach gives us far more than point estimates of preference. First, it addresses the uncertainty embedded in observational data and the inference process, propagating it to every derived quantity. Second, the hierarchical machinery does not stop at the preferences: we reconstruct the interaction rate of the underlying stochastic process rather than relying on a corresponding noisy sample, so every inferred variable carries ecological signal worth interpreting. Finally, the framework is flexible: from a base case for single-observation data, it extends naturally to data sampled across multiple sites, over repeated time points, and with covariates. We illustrated the framework using plant-pollinator data, but it naturally applies to other mutualistic interactions.

We first examined the model in a synthetic setting, where the underlying preferences are known and recovery can be evaluated directly. A simulated example showed how the same latent preferences can generate markedly different count patterns depending on how the system is observed and on species abundances. Raw counts therefore cannot be read as preferences on their own, and undoing exactly this confound is what our model is built to do. The effect is more pronounced in a multi-site simulated dataset, where simply aggregating counts across samples tracks abundance and fails to recover preferences, whereas our fitting procedure does. This demonstrates the need for such a framework.

Across the range of scenarios we examined, what limits recovery of the preferences is the data rather than the model. Sweeping over preference and abundance heterogeneity, count baseline, and community size, the model recovered the quantities of interest throughout, degrading only mildly toward the extremes of each regime, evidence that its performance does not hinge on a narrow, favourable corner of parameter space. What most limits our ability to separate preferences from observation is sampling effort first, and the variability of individual species’ activity second.

The multi-site results strengthen this lesson: distributing sampling across sites substantially improves reconstruction of the latent process relative to single-site sampling, especially at a lower per-site baseline and when species activity is most variable. This motivates sampling richer, more granular data of plant-pollinator communities [28].

On empirical data, the model shows how the pattern of mutualistic preferences departs from observed counts once individual species effects are removed. This effect is even more pronounced in a longitudinal, spatially resolved dataset. This is crucial given the widespread practice of relying on single-snapshot data, or aggregating finer-grained sampling into one observation [2]. Assessing species-pair relationships from summed field visits misrepresents the latent affinities, undermining any network analysis built on them. On the multi-observation data of [25], the hierarchical structure lets us quantify variability across species in sites, time, and individual activity, structure that a single fitted network cannot expose.

On the dataset of [15], adding a single binary covariate for low versus high urbanisation reveals how this acts on the community. The data support no change in baseline activity, but the effect is more pronounced for plants than pollinators, recovering, from visit counts alone, the phenological mismatch inferred indirectly by [15]. We find a (mild) signal of rewiring in mutualistic preferences, and interaction diversity separates the two urbanisation levels. Crucially, this contrast is only visible at the level of the latent preferences and not on the raw observed counts. Decomposing the dissimilarity, following [33], strengthens this picture: on observed counts the difference looks almost entirely like rewiring, but on the abundance-free preferences the rewiring contribution is markedly lower, and species turnover dominates, suggesting most of the apparent rewiring is an artefact of shifting abundances rather than a genuine change in who prefers whom. This resolves a distinction that [15] leave open, leading to a different conclusion. The implication is methodological as much as ecological. Neither extreme—aggregating counts into one snapshot, nor treating each site–time sample as an isolated network—recovers these signals, which emerge only by borrowing strength across observations to infer the shared latent process, otherwise masked by the noise of any single realisation.

Like any model, the framework rests on assumptions and has limitations, which we discuss here alongside possible extensions.

The first assumption is the normal prior on the pairwise preferences *π*_*ij*_, which treats species pairs as *a priori* exchangeable: no pair is expected to share a stronger mutualistic preference than any other. Yet preferences are not arbitrary, traits and phylogenetic relationships are known to shape which species associate [10, 35]. The exchangeable prior does not deny this, it simply lets such structure emerge from the data, rather than wiring it in from the start. A natural extension would encode that structure directly, through a phylogenetically correlated prior or trait-based matching.

A second assumption concerns the likelihood. As is standard for ecological count data, we adopt a Poisson observation model, in which mean and variance coincide. Interaction counts, however, can be overdispersed [5], and where they are, the model extends easily to a negative-binomial alternative.

A more consequential limitation is that observed data are almost always incomplete, with some interactions going un-recorded within the sampling window [23]. Our model takes the observed visits at face value: with no detection or zero-inflation layer, every unseen interaction enters the likelihood as a genuine Poisson zero, that pushes the expected rate downward. In practice, distinguishing whether that low rate reflects a low abundance baseline or a genuine avoidance is a pitfall only in specific circumstances. The quantity we target is the preference that remains once the abundance baseline has been removed, and whether a zero informs it depends entirely on how well that baseline is pinned down. When both species are well sampled and their baseline predicts a non-trivial number of visits, an unseen interaction falls well below that expectation and is correctly read as evidence of *low* preference. The genuine exposure lies with rare species seen interacting with only one or two partners, where the baseline is poorly determined and a single unrecorded interaction can leave a real preference with no support in the data.

Modelling the observation process explicitly is a natural next step, though at a real cost in complexity. [1] separate a latent ecological network from an observation layer driven by abundance and sampling, but their latent object is a *binary* network, with no pair-specific preference. [49] likewise model imperfect detection as a binomial thinning of a latent Poisson intensity, though aimed at low-rank connectivity rather than heterogeneous preferences. Bringing an explicit missingness model together with pair-specific preference estimation is, to our knowledge, still open.

More generally, a Bayesian treatment reports only what the data support: the posterior stays broad, close to the pooled prior for rarely seen pairs, and concentrates for well-sampled ones. This is a point of careful interpretation rather than a limitation. A wide posterior signals that, once confounding effects are removed, the data do not justify a strong preference. Partial pooling handles this coherently, borrowing strength where a single pair is uninformative, though sharp conclusions still require adequately sampled data.

Together, these results show the importance of treating observed plant-pollinator visits as the starting point for inferring the latent process, not the final object of analysis. By separating, in a Bayesian setting, mutualistic preference from abundance, sampling, and site- and time-specific effects, our framework recovers a signal that raw counts obscure. This speaks to a growing recognition of the value of such reconstruction; because it acts on the foundational step of any network analysis, its implications are wide, representing a further step towards more grounded network-based approaches in ecology.

## DATA AND CODE AVAILABILITY

We fit the model in Stan [43] via `cmdstanpy` [42], with full sampler settings and convergence diagnostics in the SI (Sec. S1). All code required to reproduce the analyses and simulations in this study, including the fully reproducible synthetic datasets, is available at https://github.com/leonardofederici/DELPHI.

The single-observation empirical network, as well as the additional datasets used in the supplementary analyses, are taken from the compilation of previously published pollination networks provided in the supplementary information of [36]. The restoration data of

[25] are available from the Interaction Web DataBase (http://www.ecologia.ib.usp.br/iwdb/html/kaiser-bunbury_et_al_2017.html). The seasonal urbanisation data of [15] are available from Zenodo (https://doi.org/10.5281/zenodo.5570297). Full citations for all source publications are given in the reference list.

## AUTHOR CONTRIBUTIONS

All authors conceived and designed the study. L.F. performed all analyses and wrote the first draft of the manuscript. E.M. and I.I. jointly supervised the project. All authors reviewed and edited the manuscript and approved the final version.

## References

[1] E. Anakok, P. Barbillon, C. Fontaine, and E. Thebault. Disentangling the structure of ecological bipartite networks from observation processes. The Annals of Applied Statistics, 19(1): 397–418, 2025.

[2] J. Bascompte. Web of life: ecological networks database, 2023. URL http://www.web-of-life.es.

[3] J. Bascompte, P. Jordano, C. J. Melián, and J. M. Olesen. The nested assembly of plant–animal mutualistic networks. Proceedings of the National Academy of Sciences, 100(16):9383–9387, 2003.

[4] U. Bastolla, M. A. Fortuna, A. Pascual-García, A. Ferrera, B. Luque, and J. Bascompte. The architecture of mutualistic networks minimizes competition and increases biodiversity. Nature, 458(7241):1018–1020, 2009. ISSN 1476-4687.

[5] C. I. Bliss and R. A. Fisher. Fitting the negative binomial distribution to biological data. Biometrics, 9(2):176–200, 1953.

[6] N. Blüthgen and M. Staab. A critical evaluation of network approaches for studying species interactions. Annual Review of Ecology, Evolution, and Systematics, 55(1):65–88, 2024.

[7] N. Blüthgen, F. Menzel, and N. Blüthgen. Measuring specialization in species interaction networks. BMC ecology, 6(1):9, 2006.

[8] J. L. Bronstein, editor. Mutualism. Oxford University Press, 2016.

[9] P. J. CaraDonna, L. A. Burkle, B. Schwarz, J. Resasco, T. M. Knight, G. Benadi, N. Blüthgen, C. F. Dormann, Q. Fang, J. Fründ, et al. Seeing through the static: the temporal dimension of plant–animal mutualistic interactions. Ecology Letters, 24(1): 149–161, 2021.

[10] S. A. Chamberlain, R. V. Cartar, A. C. Worley, S. J. Semmler, G. Gielens, S. Elwell, M. E. Evans, J. C. Vamosi, and E. Elle. Traits and phylogenetic history contribute to network structure across canadian plant–pollinator communities. Oecologia, 176 (2):545–556, 2014.

[11] D. De Moor, J. D. Hart, D. W. Franks, L. J. Brent, M. J. Silk, and J. B. Brask. Latent layers in social networks and their implications for comparative analyses. Behavioral Ecology, 36 (6):araf113, 2025.

[12] E. Delmas, M. Besson, M.-H. Brice, L. A. Burkle, G. V. Dalla Riva, M.-J. Fortin, D. Gravel, P. R. Guimarães Jr, D. H. Hembry, E. A. Newman, et al. Analysing ecological networks of species interactions. Biological Reviews, 94(1):16–36, 2019.

[13] V. Domínguez-Garcia, F. P. Molina, O. Godoy, and I. Bartomeus. Interaction network structure explains species’ temporal persistence in empirical plant–pollinator communities. Nature Ecology & Evolution, 8:423–429, 2024.

[14] A. Fisogni, N. Hautekeète, Y. Piquot, M. Brun, C. Vanappel-ghem, D. Michez, and F. Massol. Urbanization drives an early spring for plants but not for pollinators. Oikos, 129(11):1681–1691, 2020.

[15] A. Fisogni, N. Hautekeète, Y. Piquot, M. Brun, C. Vanappel-ghem, M. Ohlmann, M. Franchomme, C. Hinnewinkel, and F. Massol. Seasonal trajectories of plant-pollinator interaction networks differ following phenological mismatches along an urbanization gradient. Landscape and Urban Planning, 226: 104512, 2022.

[16] D. W. Franks, M. N. Weiss, M. J. Silk, R. J. Perryman, and D. P. Croft. Bison: A bayesian framework for inference of social networks. Methods in Ecology and Evolution, 12(9):1740–1750, 2021.

[17] J. Freimuth, O. Bossdorf, J. F. Scheepens, and F. M. Willems. Climate warming changes synchrony of plants and pollinators. Proceedings of the Royal Society B: Biological Sciences, 289 (1971), 2022.

[18] K. J. Gaston, T. M. Blackburn, J. J. Greenwood, R. D. Gregory, R. M. Quinn, and J. H. Lawton. Abundance–occupancy relationships. Journal of Applied Ecology, 37(1):39–59, 2000. doi:10.1046/j.1365-2664.2000.00485.x.

[19] A. Gelman, A. Vehtari, D. Simpson, C. C. Margossian, B. Carpenter, Y. Yao, L. Kennedy, J. Gabry, P.-C. Bürkner, and M. Modrák. Bayesian workflow. arXiv preprint arXiv:2011.01808, 2020.

[20] R. Gibson, B. Knott, T. Eberlein, and J. Memmott. Sampling method influences the structure of plant–pollinator networks. Oikos, 120, 2011.

[21] T. Harrison and R. Winfree. Urban drivers of plant-pollinator interactions. Functional Ecology, 29(7):879–888, 2015.

[22] S. Hervías-Parejo, P. Colom, R. Beltran Mas, P. E. Serra, S. Pons, V. Mesquida, and A. Traveset. Spatio-temporal variation in plant– pollinator interactions: a multilayer network approach. Oikos, 2023(8):e09818, 2023.

[23] P. Jordano. Sampling networks of ecological interactions. Functional Ecology, 30(12):1883–1893, 2016.

[24] P. Jordano, J. Bascompte, and J. M. Olesen. Invariant properties in coevolutionary networks of plant-animal interactions. Ecology Letters, 6(1):69–81, 2003.

[25] C. N. Kaiser-Bunbury, J. Mougal, A. E. Whittington, T. Valentin, R. Gabriel, J. M. Olesen, and N. Blüthgen. Ecosystem restoration strengthens pollination network resilience and function. Nature, 542(7640):223–227, 2017.

[26] N. P. Lemoine. Moving beyond noninformative priors: why and how to choose weakly informative priors in bayesian analyses. Oikos, 128(7):912–928, 2019.

[27] H. Liang, Y.-D. He, P. Theodorou, and C.-F. Yang. The effects of urbanization on pollinators and pollination: A meta-analysis. Ecology Letters, 26(9):1629–1642, 2023.

[28] I. Manning, L. Zoller, and J. Resasco. Impacts of sampling effort on seasonal plant-pollinator interaction turnover over eight years. Oecologia, 207(8):131, 2025.

[29] B. J. McGill. A test of the unified neutral theory of biodiversity. Nature, 422(6934):881–885, 2003. ISSN 1476-4687. doi:10.1038/nature01583.

[30] R. J. Mitchell, R. E. Irwin, R. J. Flanagan, and J. D. Karron. Ecology and evolution of plant–pollinator interactions. Annals of Botany, 103(9):1355–1363, 2009.

[31] M. Ohlmann, V. Miele, S. Dray, L. Chalmandrier, L. O’connor, and W. Thuiller. Diversity indices for ecological networks: a unifying framework using hill numbers. Ecology letters, 22(4): 737–747, 2019.

[32] M. Pascual and J. A. Dunne. Ecological networks: linking structure to dynamics in food webs. Oxford University Press, 2005.

[33] T. Poisot, E. Canard, D. Mouillot, N. Mouquet, and D. Gravel. The dissimilarity of species interaction networks. Ecology letters, 15(12):1353–1361, 2012.

[34] T. Poisot, D. B. Stouffer, and D. Gravel. Beyond species: why ecological interaction networks vary through space and time. Oikos, 124(3):243–251, 2015.

[35] N. E. Rafferty and A. R. Ives. Phylogenetic trait-based analyses of ecological networks. Ecology, 94(10):2321–2333, 2013.

[36] E. L. Rezende, J. E. Lavabre, P. R. Guimarães, P. Jordano, and J. Bascompte. Non-random coextinctions in phylogenetically structured mutualistic networks. Nature, 448, 2007.

[37] A. Rivera-Hutinel, R. Bustamante, V. Marín, and R. Medel. Effects of sampling completeness on the structure of plant– pollinator networks. Ecology, 93(7):1593–1603, 2012.

[38] C. Robertson. Flowers and insects: Lists of visitors of four hundred and fifty-three flowers. Science Press, 1928.

[39] R. P. Rohr, S. Saavedra, and J. Bascompte. On the structural stability of mutualistic systems. Science, 345(6195):1253497, 2014. doi:10.1126/science.1253497.

[40] S. Saavedra, D. B. Stouffer, B. Uzzi, and J. Bascompte. Strong contributors to network persistence are the most vulnerable to extinction. Nature, 478(7368):233–235, 2011.

[41] J. D. H. Sprayberry, T.-L. Ashman, J. Crall, J. Hranitz, M. Jankauski, M. Lihoreau, S. Potdar, N. E. Rafferty, C. C. Rittschof, M. A. Y. Smith, I. Valdes, and E. L. Westerman. Plant-pollinator interactions in the anthropocene: Why we need a systems approach. Integrative and Comparative Biology, page icaf062, 2025.

[42] Stan Development Team. CmdStanPy: A Python interface to CmdStan. https://github.com/stan-dev/cmdstanpy, 2024. Python package version 1.2.5.

[43] Stan Development Team. Stan Reference Manual, Version 2.36, 2025. URL https://mc-stan.org.

[44] P. P. Staniczenko, J. C. Kopp, and S. Allesina. The ghost of nestedness in ecological networks. Nature communications, 4 (1):1391, 2013.

[45] E. Thébault and C. Fontaine. Stability of ecological communities and the architecture of mutualistic and trophic networks. Science, 329(5993):853–856, 2010. ISSN 0036-8075, 1095-9203. doi: 10.1126/science.1188321.

[46] T. van der Niet, R. Peakall, and S. D. Johnson. Pollinator-driven ecological speciation in plants: new evidence and future perspectives. Annals of Botany, 113(2):199–212, 2014.

[47] D. P. Vázquez, N. Blüthgen, L. Cagnolo, and N. P. Chacoff. Uniting pattern and process in plant–animal mutualistic networks: a review. Annals of Botany, 103(9):1445–1457, 2009.

[48] J.-G. Young, F. S. Valdovinos, and M. E. Newman. Reconstruction of plant-pollinator networks from observational data. Nature Communications, 12(1):3911, 2021.

[49] A. Zhang, T. Wei, M. J. Guerrero, and C. A. Uribe. Sparse network inference under imperfect detection and its application to ecological networks. arXiv preprint arXiv:2604.18820, 2026.

