## Supplementary Information for "Inferring the latent network of pairwise mutualistic preferences from observed plant–pollinator interactions"

### CONTENTS

S1. Implementation and inference details 2

S2. Extending the model with measured covariates 3

S3. Likelihood and posterior derivation for the single-observation model 4

S4. Simulated data examples 6
- A. Convergence diagnostics 6
- B. Metric comparison of the two single-observation scenarios 6

S5. Prior predictive check 7

S6. Simulation-Based Calibration 9
- A. Setup for the current model 9
  - Results 10

S7. Prior robustness 12

S8. Model recovery performance 14
- A. Single-observation model 14
  - Results 15
- B. Multi-site model 15
  - Results 16

S9. Single-observation empirical case study 22

S10. Multi-observation empirical case study 23

S11. Urbanisation gradient case study 24
- A. Full model description 24
- B. Posterior summaries 25
- C. Convergence diagnostics 25
- D. Network metrics 25
- E. Interaction $\beta$-diversity decomposition 26

References 28

### S1. IMPLEMENTATION AND INFERENCE DETAILS

All results were produced with CmdStan 2.36 and `cmdstanpy` 1.2.5 under Python 3.13. Posterior sampling used Stan's adaptive Hamiltonian Monte Carlo, specifically the no-U-turn sampler (NUTS) [7], usually with $4$ chains of $1000$ warmup and $1000$ post-warmup iterations each, increased to 1500 or 2000 for a few fits to meet convergence criteria. To accommodate the hierarchical posterior geometry, we set the target acceptance rate to $\texttt{adapt\_delta} = 0.99$ and the maximum tree depth to $15$, leaving all other sampler settings at their defaults. For every fit, we assessed convergence using the rank-normalized split-$\hat{R}$ statistic together with the bulk and tail effective sample sizes (ESS) [14]. We required $\hat{R} < 1.01$ and both bulk- and tail-ESS above $400$ for all parameters, and verified the absence of divergent transitions, no saturation of the maximum tree depth, and a satisfactory energy Bayesian fraction of missing information (E-BFMI).

### S2. EXTENDING THE MODEL WITH MEASURED COVARIATES

If the dataset includes measured spatial or temporal covariates, such as habitat type or temperature, these can be incorporated to explain part of the site- or time-level variation in species effects. We first consider the case in which the covariate acts on these species-level deviations alone, and generalise to other components of the model at the end of this section. Rather than treating sites and time points as exchangeable, we split the corresponding deviations into a component explained by the covariate and a residual:

$$\begin{aligned}\delta^p_{i,s} &= \alpha^p_i\, x_s + \epsilon^p_{i,s},\\ \eta^p_{i,t} &= \beta^p_i\, z_t + \xi^p_{i,t},\end{aligned} \tag{S1}$$

where $x_s$ and $z_t$ are site- and time-level covariates, the species-specific slopes $\alpha^p_i$, $\beta^p_i$ quantify how plant species $i$ responds to each, and the residuals $\epsilon^p_{i,s}$, $\xi^p_{i,t}$ capture what remains unexplained. The same structure applies to pollinator effects $a_j^{(s,t)}$ and to the intercept $\gamma^{(s,t)}$.

Following the same hierarchical logic, we assign random-effects priors to slopes and residuals:

$$\begin{aligned}\alpha^p_i &\sim \mathcal{N}(0, \sigma^2_\alpha), & \epsilon^p_{i,s} &\sim \mathcal{N}(0, {\Sigma^\epsilon_p}^2),\\ \beta^p_i &\sim \mathcal{N}(0, \sigma^2_\beta), & \xi^p_{i,t} &\sim \mathcal{N}(0, {\Sigma^\xi_p}^2).\end{aligned} \tag{S2}$$

The zero-centred priors on $\alpha^p_i$ and $\beta^p_i$ express the assumption that species responses to the covariate are exchangeable: before seeing data, no species is expected to respond more strongly than any other. Here $\sigma_\alpha$ ($\sigma_\beta$) measures how much species differ in their response to the spatial (temporal) covariate, while the zero-centred slopes leave any average community-wide response to be absorbed by the intercept.

The residual variances $\Sigma^\epsilon_p$ and $\Sigma^\xi_p$ play the role previously held by $\Sigma_p$ and $\Sigma_{p,t}$, but now capture only the variation *not* explained by the covariates. A reduction of $\Sigma^\epsilon_p$ relative to $\Sigma_p$ indicates that the covariate accounts for some of the between-site variation.

The extension to $K$ covariates is straightforward, replacing each slope term with a sum $\sum_{k=1}^{K} \alpha^p_{i,k}\, x_{s,k}$ whose slopes each receive their own hierarchical prior $\alpha^p_{i,k} \sim \mathcal{N}(0, \sigma^2_{\alpha_k})$, and analogously in time. In all cases, the modular structure of the random-effects framework ensures that covariates can be added or removed without altering the rest of the model architecture. The introduction of $K$ covariates increases model complexity by $K \times (N_p + N_a)$ slope parameters, though the hierarchical priors on the slopes control the effective dimensionality by shrinking poorly informed responses toward zero. In practice, the number of covariates should remain modest relative to the number of sites (or time points), to ensure that the covariate structure captures generalisable ecological responses rather than overfitting site-level variation.

So far we have let the covariate act on the site- and time-level deviations of the species effects alone, holding the mutualistic preferences $\pi_{ij}$ invariant. More generally, depending on its nature and on the question of interest, the same linear-predictor construction applies to other channels of the model, such as the baseline and the preferences themselves. This is the route taken in S11, where the covariate defines a contrast between two conditions and is therefore allowed to modulate the preferences as well.

### S3. LIKELIHOOD AND POSTERIOR DERIVATION FOR THE SINGLE-OBSERVATION MODEL

Before deriving the joint posterior of the single-observation model of the main text, we first recall the model hierarchical structure. We start from observed visit counts matrix $\mathbf{M} = \{M_{ij}\}$, where $i = 1, \dots, n_\mathrm{p}$ indexes plants and $j = 1, \dots, n_\mathrm{a}$ indexes pollinators. Each count is modelled as a Poisson realisation whose rate is decomposed on the log scale,

$$M_{ij} \mid \mu_{ij} \sim \mathrm{Poisson}(\mu_{ij}), \qquad \log \mu_{ij} = \gamma + p_i + a_j + \pi_{ij}.$$

The species effects and the pairwise preferences are assigned zero-mean Gaussian priors,

$$\gamma \mid \sigma_\gamma \sim \mathcal{N}(0, \sigma_\gamma^2), \qquad p_i \mid \sigma_p \sim \mathcal{N}(0, \sigma_p^2), \qquad a_j \mid \sigma_a \sim \mathcal{N}(0, \sigma_a^2), \qquad \pi_{ij} \mid \sigma_\pi \sim \mathcal{N}(0, \sigma_\pi^2),$$

and the scale parameters are given exponential hyperpriors,

$$\sigma_\gamma \sim \mathrm{Exp}(\lambda_\gamma), \qquad \sigma_p, \sigma_a, \sigma_\pi \sim \mathrm{Exp}(\lambda). \tag{S3}$$

In the main text $\sigma_\gamma$ is held fixed at a weakly informative value. Here we consider $\sigma_\gamma$ as an additional stochastic node with its own hyperprior in Eq. (S3). The case of fixed $\sigma_\gamma$ is recovered by removing the terms in $\sigma_\gamma$ from the hyperprior and reading it as a constant. We group the model parameters into main parameters and hyper-parameters,

$$\boldsymbol{\theta} = \big(\gamma,\ \{p_i\}_{i=1}^{n_\mathrm{p}},\ \{a_j\}_{j=1}^{n_\mathrm{a}},\ \{\pi_{ij}\}_{i,j}\big), \qquad \boldsymbol{\phi} = \big(\sigma_\gamma,\ \sigma_p,\ \sigma_a,\ \sigma_\pi\big).$$

By Bayes' theorem the joint posterior of the parameters is

$$P(\boldsymbol{\theta}, \boldsymbol{\phi} \mid \mathbf{M}) = \frac{P(\mathbf{M} \mid \boldsymbol{\theta}, \boldsymbol{\phi})\, P(\boldsymbol{\theta}, \boldsymbol{\phi})}{P(\mathbf{M})}, \qquad P(\mathbf{M}) = \iint P(\mathbf{M}, \boldsymbol{\theta}, \boldsymbol{\phi})\, \mathrm{d}\boldsymbol{\theta}\, \mathrm{d}\boldsymbol{\phi}.$$

The observation model does not depend on the hyper-parameters once the main parameters are fixed, so $P(\mathbf{M} \mid \boldsymbol{\theta}, \boldsymbol{\phi}) = P(\mathbf{M} \mid \boldsymbol{\theta})$, that is $\mathbf{M} \perp \boldsymbol{\phi} \mid \boldsymbol{\theta}$. The prior factorises as $P(\boldsymbol{\theta}, \boldsymbol{\phi}) = P(\boldsymbol{\theta} \mid \boldsymbol{\phi})\, P(\boldsymbol{\phi})$ by the definition of conditional probability. The evidence $P(\mathbf{M})$ is a high-dimensional integral with no closed form. It is also not required for posterior sampling, because Markov chain Monte Carlo methods use the posterior only through density ratios or through the gradient of its logarithm, in both of which the constant $P(\mathbf{M})$ cancels. We therefore work with the posterior up to proportionality,

$$P(\boldsymbol{\theta}, \boldsymbol{\phi} \mid \mathbf{M}) \ \propto\ P(\mathbf{M} \mid \boldsymbol{\theta})\, P(\boldsymbol{\theta} \mid \boldsymbol{\phi})\, P(\boldsymbol{\phi}). \tag{S4}$$

*Likelihood.* Conditional on the rates, the pairwise visit counts are considered independent across cells, so the likelihood is a product of Poisson terms,

$$P(\mathbf{M} \mid \boldsymbol{\theta}) = \prod_{i=1}^{n_\mathrm{p}} \prod_{j=1}^{n_\mathrm{a}} \mathrm{Poisson}(M_{ij}; \mu_{ij}) = \prod_{i=1}^{n_\mathrm{p}} \prod_{j=1}^{n_\mathrm{a}} \frac{\mu_{ij}^{M_{ij}}\, e^{-\mu_{ij}}}{M_{ij}!}.$$

Taking logarithms gives the log-likelihood

$$\ell(\boldsymbol{\theta}) := \log P(\mathbf{M} \mid \boldsymbol{\theta}) = \sum_{i,j} \big[\, M_{ij} \log \mu_{ij} - \mu_{ij} - \log(M_{ij}!) \,\big].$$

Substituting the log-rate $\log \mu_{ij} = \gamma + p_i + a_j + \pi_{ij}$ yields

$$\ell(\boldsymbol{\theta}) = \sum_{i,j} \big[\, M_{ij}(\gamma + p_i + a_j + \pi_{ij}) - \exp(\gamma + p_i + a_j + \pi_{ij}) \,\big] - \sum_{i,j} \log(M_{ij}!).$$

Expanding the linear term and collecting by parameter gives

$$\sum_{i,j} M_{ij} \log \mu_{ij} = \gamma\, M_{\bullet\bullet} + \sum_i p_i\, M_{i\bullet} + \sum_j a_j\, M_{\bullet j} + \sum_{i,j} \pi_{ij} M_{ij},$$

with grand total, row totals and column totals

$$M_{\bullet\bullet} = \sum_{i,j} M_{ij}, \qquad M_{i\bullet} = \sum_j M_{ij}, \qquad M_{\bullet j} = \sum_i M_{ij}. \tag{S5}$$

Each plant effect $p_i$ enters the likelihood only through its row total $M_{i\bullet}$, and each pollinator effect $a_j$ only through its column total $M_{\bullet j}$, whereas each preference $\pi_{ij}$ is informed by the single count $M_{ij}$.

*Priors and hyperpriors.* Given the hyper-parameters, the components of $\boldsymbol{\theta}$ are independent, so the prior factorises across the four blocks,

$$P(\boldsymbol{\theta} \mid \boldsymbol{\phi}) = \mathcal{N}(\gamma; 0, \sigma_\gamma^2) \prod_{i=1}^{n_\mathrm{p}} \mathcal{N}(p_i; 0, \sigma_p^2) \prod_{j=1}^{n_\mathrm{a}} \mathcal{N}(a_j; 0, \sigma_a^2) \prod_{i,j} \mathcal{N}(\pi_{ij}; 0, \sigma_\pi^2).$$

Writing out the Gaussian densities and collecting terms,

$$\begin{aligned} P(\boldsymbol{\theta} \mid \boldsymbol{\phi}) = {} & (2\pi)^{-\frac{1+n_\mathrm{p}+n_\mathrm{a}+n_\mathrm{p}n_\mathrm{a}}{2}} \, \sigma_\gamma^{-1} \, \sigma_p^{-n_\mathrm{p}} \, \sigma_a^{-n_\mathrm{a}} \, \sigma_\pi^{-n_\mathrm{p}n_\mathrm{a}} \\ & \times \exp\left( -\frac{\gamma^2}{2\sigma_\gamma^2} - \frac{1}{2\sigma_p^2} \sum_i p_i^2 - \frac{1}{2\sigma_a^2} \sum_j a_j^2 - \frac{1}{2\sigma_\pi^2} \sum_{i,j} \pi_{ij}^2 \right), \end{aligned}$$

so that

$$\begin{aligned} \log P(\boldsymbol{\theta} \mid \boldsymbol{\phi}) = {} & -\tfrac{1+n_\mathrm{p}+n_\mathrm{a}+n_\mathrm{p}n_\mathrm{a}}{2} \log(2\pi) - \log \sigma_\gamma - n_\mathrm{p} \log \sigma_p - n_\mathrm{a} \log \sigma_a - n_\mathrm{p} n_\mathrm{a} \log \sigma_\pi \\ & - \frac{\gamma^2}{2\sigma_\gamma^2} - \frac{1}{2\sigma_p^2} \sum_i p_i^2 - \frac{1}{2\sigma_a^2} \sum_j a_j^2 - \frac{1}{2\sigma_\pi^2} \sum_{i,j} \pi_{ij}^2. \end{aligned}$$

The exponential hyperpriors, $\mathrm{Exp}(x; \lambda) = \lambda e^{-\lambda x}$ for $x > 0$, contribute

$$P(\boldsymbol{\phi}) = \mathrm{Exp}(\sigma_\gamma; \lambda_\gamma) \, \mathrm{Exp}(\sigma_p; \lambda) \, \mathrm{Exp}(\sigma_a; \lambda) \, \mathrm{Exp}(\sigma_\pi; \lambda),$$

with logarithm

$$\log P(\boldsymbol{\phi}) = \log \lambda_\gamma + 3 \log \lambda - \lambda_\gamma \, \sigma_\gamma - \lambda(\sigma_p + \sigma_a + \sigma_\pi).$$

*Log-posterior.* Combining the likelihood, the prior and the hyperprior through Eq. (S4) and taking logarithms gives the log-posterior

$$\begin{aligned} \log P(\boldsymbol{\theta}, \boldsymbol{\phi} \mid \mathbf{M}) = {} & \sum_{i,j} \big[ M_{ij}(\gamma + p_i + a_j + \pi_{ij}) - \exp(\gamma + p_i + a_j + \pi_{ij}) \big] \\ & - \frac{\gamma^2}{2\sigma_\gamma^2} - \frac{1}{2\sigma_p^2} \sum_i p_i^2 - \frac{1}{2\sigma_a^2} \sum_j a_j^2 - \frac{1}{2\sigma_\pi^2} \sum_{i,j} \pi_{ij}^2 \\ & - \log \sigma_\gamma - n_\mathrm{p} \log \sigma_p - n_\mathrm{a} \log \sigma_a - n_\mathrm{p} n_\mathrm{a} \log \sigma_\pi \\ & - \lambda_\gamma \, \sigma_\gamma - \lambda(\sigma_p + \sigma_a + \sigma_\pi) + \mathrm{const}, \end{aligned}$$

where the additive constant collects all terms independent of $(\boldsymbol{\theta}, \boldsymbol{\phi})$, namely $-\log P(\mathbf{M})$, $-\sum_{ij} \log(M_{ij}!)$, $-\frac{1}{2}(1 + n_\mathrm{p} + n_\mathrm{a} + n_\mathrm{p} n_\mathrm{a}) \log(2\pi)$ and $\log \lambda_\gamma + 3 \log \lambda$.

Using the totals defined in Eq. (S5) and writing $\mu_{\bullet\bullet} = \sum_{ij} \mu_{ij} = \sum_{ij} \exp(\gamma + p_i + a_j + \pi_{ij})$ for the total rate, the log-posterior can be written equivalently as

$$\begin{aligned} \log P(\boldsymbol{\theta}, \boldsymbol{\phi} \mid \mathbf{M}) = {} & \gamma N + \sum_i p_i \, M_{i\bullet} + \sum_j a_j \, M_{\bullet j} + \sum_{i,j} \pi_{ij} M_{ij} - \mu_{\bullet\bullet} \\ & - \frac{\gamma^2}{2\sigma_\gamma^2} - \frac{1}{2\sigma_p^2} \sum_i p_i^2 - \frac{1}{2\sigma_a^2} \sum_j a_j^2 - \frac{1}{2\sigma_\pi^2} \sum_{i,j} \pi_{ij}^2 \\ & - \log \sigma_\gamma - n_\mathrm{p} \log \sigma_p - n_\mathrm{a} \log \sigma_a - n_\mathrm{p} n_\mathrm{a} \log \sigma_\pi - \lambda_\gamma \, \sigma_\gamma - \lambda(\sigma_p + \sigma_a + \sigma_\pi) + \mathrm{const}. \end{aligned}$$

In this form the data enter the parameter-dependent part only through the summary statistics $M_{\bullet\bullet}$, $M_{i\bullet}$ and $M_{\bullet j}$, the individual counts $M_{ij}$, and the total rate $\mu_{\bullet\bullet}$. The species effects are thus informed by the corresponding row and column totals and each preference by a single count, in line with the partial pooling induced by the hierarchical priors.

### S4. SIMULATED DATA EXAMPLES

#### A. Convergence diagnostics

Across the three synthetic settings considered in the main text—namely, Scenarios I and II of the single-observation case (Fig. 2) and the multi-site data (Fig. 3)—we assessed the convergence of the sampler using the criteria described above. For every fit, the rank-normalized split-$\hat{R}$ remained below $1.01$ for all parameters, and no fit produced divergent transitions, saturated the maximum tree depth, or returned an unsatisfactory energy diagnostic (E-BFMI). Both bulk- and tail-effective sample sizes exceeded the target of $400$ for all parameters in every setting, the smallest per-setting values being a bulk-ESS of $870$, $1045$, and $1014$ and a tail-ESS of $851$, $1622$, and $1744$, respectively. We therefore regard the posterior summaries reported for the synthetic examples as reflecting adequate sampling of the posterior.

#### B. Metric comparison of the two single-observation scenarios

To show that relying on observed counts can produce different pictures even when the underlying preferences are identical, we compare the two scenarios presented in Fig. 2 using two complementary measures: the relative Frobenius distance and the Spearman rank correlation. The first quantifies agreement between two matrices in terms of magnitude, while the second disregards absolute magnitudes and captures instead the agreement in the ranking of the entries.

For two matrices $X$ and $Y$, the Frobenius norm is $\|X\|_F = \big(\sum_{i,j} X_{ij}^2\big)^{1/2}$, and we define the relative Frobenius distance as

$$d_F(X, Y) = \frac{\|X - Y\|_F}{\frac{1}{2}\big(\|X\|_F + \|Y\|_F\big)},$$

which normalises the discrepancy by the typical magnitude of the two matrices and is therefore comparable across quantities measured on different scales. The Spearman correlation $\rho$ is computed between the vectorised entries of the two matrices, that is, on the ranks of the corresponding pairs.

Applying these to the two observed count matrices yields a relative Frobenius distance of $d_F = 1.73$ and a Spearman correlation of $\rho = 0.66$. Repeating the analysis on the inferred preference matrices, with each entry taken as its posterior median, yields $d_F = 0.32$ and $\rho = 0.95$.

Because the Frobenius distance is normalised, its values are directly comparable across the two layers. Together, the two measures show that inferring the latent preferences recovers the fact that the observed counts originate from the same underlying mutualistic structure, whereas the counts themselves diverge under the influence of species abundances and the count baseline.

### S5. PRIOR PREDICTIVE CHECK

Prior predictive checks generate synthetic data from the prior distribution alone, serving as a principled diagnostic for whether the chosen priors are compatible with domain knowledge before any data are observed [5]. Specifically, we run the model in a prior-only mode, sampling 500 times the latent parameters from their priors and pass them through the likelihood to produce, for each draw, a full synthetic count matrix, without ever conditioning on the observed visits. As shown in Fig. S1 and reported in Table S1, both visual inspection of prior-generated count matrices and summary statistics of the corresponding latent variables confirm that the selected priors are sufficiently diffuse to span a wide range of plausible scenarios, while still producing a coherent and ecologically sensible picture.

To keep the check anchored to the scale of the real problem, we repeat this for ten empirical dataset, fixing the matrix dimensions $(n_{\text{plants}}, n_{\text{polls}})$ to their observed values before drawing from the prior, and we compare each observed summary statistic against its prior predictive distribution (Table S2. Although the priors are deliberately weak and wide so that, by construction, they generate different kinds of datasets than what we observe, none of the summary statistics falls in the tails of its prior predictive distribution, indicating no serious mismatch between our prior assumptions and the observed data.

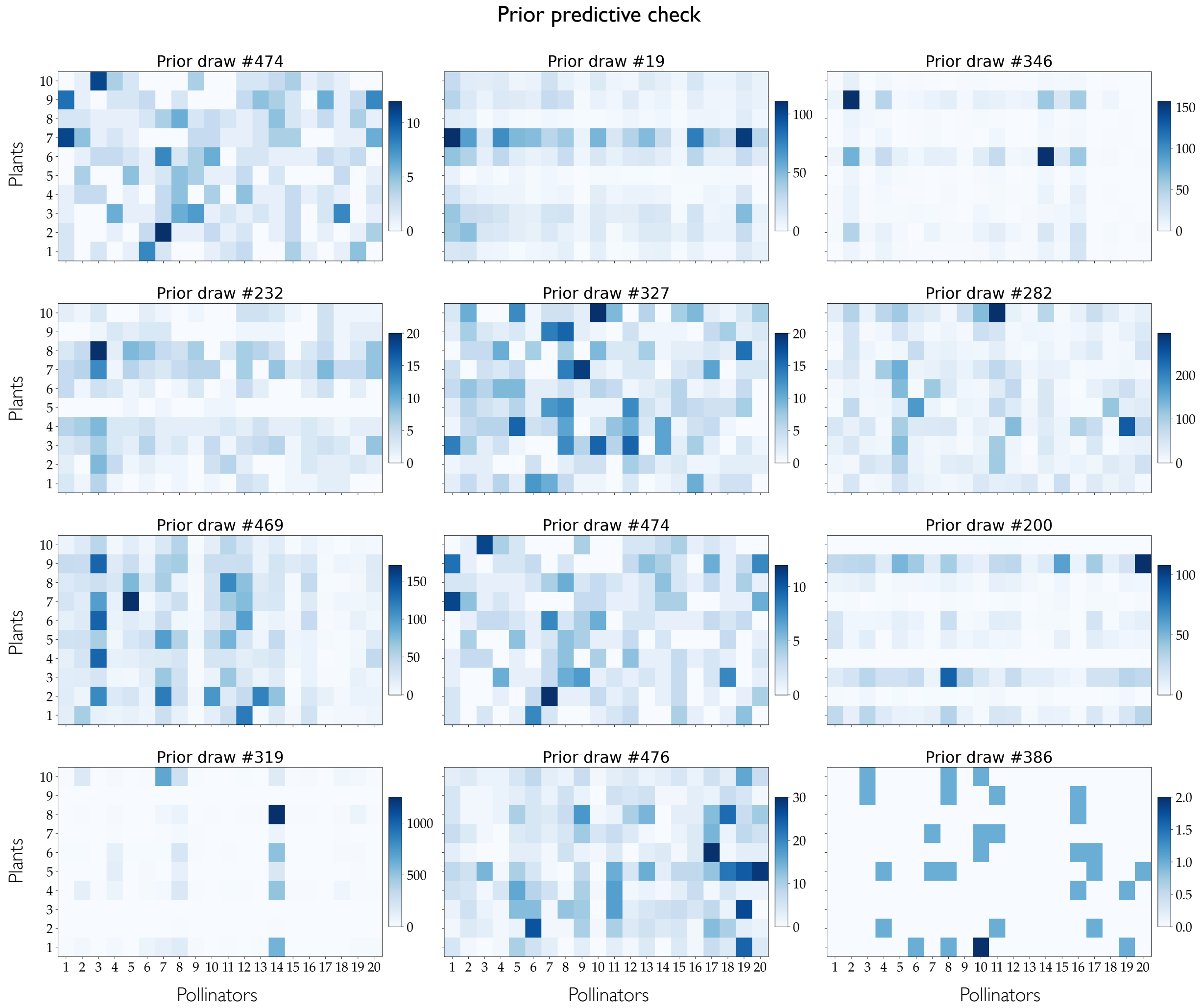


FIG. S1. **Prior predictive check.** Each matrix shows one dataset simulated from the single-observation model by drawing parameters from the prior and generating data from the likelihood, without conditioning on observed data. The spread of draws illustrates the range of outcomes the model considers plausible *a priori*, indicating that the prior encodes reasonable assumptions before fitting. Most draws fall in a sensible range, while extreme scenarios (such as the bottom-left and bottom-right heatmaps) receive lower probability.

| Parameter | Mean | Median | 5% | 95% |
|---|---|---|---|---|
| $\sigma$ | 1.303 | 1.046 | 0.893 | 2.531 |
| $\tau$ | 1.803 | 1.064 | 0.951 | 2.853 |
| $w$ | 1.146 | 1.062 | 0.999 | 1.546 |
| $\mu$ | 24.680 | 1.958 | 0.069 | 56.471 |

TABLE S1. Prior predictive check, summary statistics from prior draws of latent quantities.

| Dataset ID | | Grand total mean | Grand total SD | Grand total max | Proportion of zeros | Mean column sum | Mean row sum |
|---|---|---|---|---|---|---|---|
| | Observed | 0.45 | 3.15 | 64.00 | 0.86 | 5.39 | 45.83 |
| **BAHE** | Prior mean | 28.70 | 92.09 | 1372.53 | 0.40 | 344.38 | 2927.21 |
| | $p$-value | 0.744 | 0.446 | 0.304 | 0.096 | 0.744 | 0.744 |
| | Observed | 1.70 | 20.28 | 469.00 | 0.85 | 27.22 | 61.25 |
| **DISH** | Prior mean | 30.16 | 63.85 | 789.46 | 0.38 | 482.54 | 1085.72 |
| | $p$-value | 0.548 | 0.174 | 0.104 | 0.116 | 0.548 | 0.548 |
| | Observed | 3.04 | 35.53 | 965.00 | 0.86 | 51.64 | 185.29 |
| **DIHI** | Prior mean | 15.66 | 31.84 | 417.70 | 0.42 | 266.23 | 955.30 |
| | $p$-value | 0.412 | 0.116 | 0.066 | 0.126 | 0.412 | 0.412 |
| | Observed | 0.41 | 4.45 | 185.00 | 0.92 | 17.16 | 34.74 |
| **INPK** | Prior mean | 53.25 | 380.48 | 6922.21 | 0.43 | 2236.37 | 4525.99 |
| | $p$-value | 0.750 | 0.374 | 0.212 | 0.070 | 0.750 | 0.750 |
| | Observed | 1.11 | 7.35 | 139.00 | 0.85 | 27.63 | 87.32 |
| **MEMM** | Prior mean | 38.80 | 187.82 | 3915.60 | 0.41 | 969.99 | 3065.18 |
| | $p$-value | 0.596 | 0.266 | 0.206 | 0.126 | 0.596 | 0.596 |
| | Observed | 0.68 | 2.37 | 19.00 | 0.81 | 7.44 | 12.18 |
| **MOMA** | Prior mean | 30.66 | 53.19 | 396.50 | 0.41 | 337.25 | 551.86 |
| | $p$-value | 0.654 | 0.474 | 0.428 | 0.174 | 0.654 | 0.654 |
| | Observed | 3.89 | 27.74 | 418.00 | 0.75 | 50.57 | 171.15 |
| **MOTT** | Prior mean | 34.41 | 121.79 | 2117.58 | 0.44 | 447.33 | 1514.04 |
| | $p$-value | 0.322 | 0.130 | 0.084 | 0.238 | 0.322 | 0.322 |
| | Observed | 1.18 | 7.51 | 139.00 | 0.80 | 10.61 | 66.00 |
| **OLLE** | Prior mean | 27.20 | 72.26 | 867.44 | 0.41 | 244.83 | 1523.36 |
| | $p$-value | 0.576 | 0.282 | 0.180 | 0.186 | 0.576 | 0.576 |
| | Observed | 2.24 | 6.77 | 87.00 | 0.68 | 29.18 | 76.31 |
| **SMAL** | Prior mean | 13.91 | 27.36 | 240.87 | 0.40 | 180.82 | 472.91 |
| | $p$-value | 0.470 | 0.274 | 0.226 | 0.246 | 0.470 | 0.470 |

TABLE S2. **Prior predictive check across empirical datasets.** We consider 9 empirical datasets from the supplementary material of [11]. For each empirical dataset we draw 500 matrices from the prior-only model, fixing the dimensions $(n_{\text{plants}}, n_{\text{polls}})$ to their observed values and generating data from the likelihood without conditioning on the observed visits. For six summary statistics we report the observed value, the mean over the prior predictive draws, and a prior predictive $p$-value [2], defined as the probability of the observed statistic to be lower or equal than the one from the prior predictive distribution. The null being tested is the prior and likelihood *jointly*, so a value deep in either tail indicates prior–data conflict in the combined model. We treat the $0.05/0.95$ bounds as an heuristic diagnostic for such conflict. Crucially, we are adopting deliberately diffuse, weakly informative prior, *meant* to generate more extreme data than observed. For all six statistics and all ten datasets the $p$-values lie within $[0.05, 0.95]$, showing the observed data sit comfortably within the prior predictive support, with only the proportion of zeros for the sparsest datasets (INPK 0.070, BAHE 0.096) approaching the lower screen. Note, the three location statistics (grand total mean, mean column sum, mean row sum) share an identical $p$-value within each dataset because all three are monotone rescalings of the grand total by a fixed constant.

### S6. SIMULATION-BASED CALIBRATION

Since we are working in a generative-model setting, we assess the quality of our inference using a simulation-based calibration (SBC) check. SBC tests whether the draws returned by a posterior sampler are consistent with the model the sampler is intended to fit [3, 13]. It rests on a self-consistency property of exact Bayesian inference, that if a parameter value is drawn from the prior and data are then simulated conditional on it, that parameter value is itself a valid draw from the corresponding posterior. A correctly implemented sampler must reproduce this property, so systematic departures from it indicate a flaw in the inference. Importantly, this calibration is limited exclusively to the computational aspect of the model.

Explicitly, drawing a parameter from the prior, $\theta^{\text{sim}} \sim p(\theta)$, and then data conditional on it, $y^{\text{sim}} \sim p(y \mid \theta^{\text{sim}})$, produces a pair $(y^{\text{sim}}, \theta^{\text{sim}})$ that is, by definition, a single draw from the joint distribution $p(y, \theta)$. Since $p(\theta \mid y) \propto p(y, \theta)$, it follows that $\theta^{\text{sim}}$ is itself a draw from the posterior $p(\theta \mid y^{\text{sim}})$ induced by the data it generated. Consequently, if an algorithm samples that posterior correctly, the prior draw $\theta^{\text{sim}}$ and the algorithm's posterior draws are draws from the same distribution, and $\theta^{\text{sim}}$ should be statistically indistinguishable from them.

Concretely, one generates $N$ independent prior–data pairs $(y^{\text{sim}(n)}, \theta^{\text{sim}(n)}) \sim p(y, \theta)$ and, for each, obtains $M$ posterior draws $\theta^{(n,1)}, \ldots, \theta^{(n,M)} \sim p(\theta \mid y^{\text{sim}(n)})$ from the algorithm under test. For parameter $k$ in simulation $n$, the test quantity is the *rank* of the simulated value among the posterior draws,

$$r_{n,k} \;=\; \sum_{m=1}^{M} \mathbb{I}\left[\theta_k^{(n,m)} < \theta_k^{\text{sim}(n)}\right], \tag{S6}$$

i.e. the number of posterior draws falling below the value that generated the data. Under exact sampling these ranks take values in $\{0, \ldots, M\}$ and, by the calibration argument above, are marginally discrete-uniform, so that $r_{n,k} + 1 \sim \text{Categorical}\left(\frac{1}{M+1}, \ldots, \frac{1}{M+1}\right)$. The calibration check thus reduces to testing the collection $\{r_{n,k}\}_{n=1}^{N}$ for uniformity for each parameter $k$. Because the uniformity tests assume independent draws, the chains are thinned to approximately the effective sample size before the ranks in (S6) are computed [12].

#### A. Setup for the current model

We apply the procedure above to the single-observation model. However, monitoring every latent coordinate is unnecessary. Within each block the components share a prior and enter the likelihood symmetrically, so they are exchangeable and their marginal rank distributions coincide. For this reason, a single representative coordinate is a sufficient marginal check for the whole block. We therefore monitor one component per block, $p_1$, $a_1$, and $\pi_{11}$, together with the four top-level parameters $\gamma$, $\sigma_p$, $\sigma_a$, and $\sigma_\pi$. We also consider the parts of the posterior that couple these effects, and additionally track three cells of $\log \mu$ chosen to share no plant or pollinator index, $\log \mu_{11}$, $\log \mu_{5,10}$, and $\log \mu_{10,20}$. These composite quantities double as an implementation check, since an indexing error in the rank bookkeeping would surface most readily where several parameters are combined. This gives $K = 10$ monitored quantities. We run $N = 200$ independent simulations. Each simulation draws its own parameters and data, fits a single chain with 1000 warmup and 2000 sampling iterations , and thins by a factor of ten to $M = 200$ retained draws. The rank indicators of Eq. (S6) are computed from each fit and summed over draws to form $r_{n,k}$.

We summarize the ranks in three complementary ways. Rank histograms (Figure S2) show the empirical distribution of $\{r_{n,k}\}_n$ for each quantity against the flat profile expected under uniformity, where a bowl shape would indicate an over-concentrated posterior, a dome an over-dispersed one, and a monotone tilt a location bias. As a sharper test, we form the normalized ranks $z_{n,k} = r_{n,k}/M \in [0, 1]$, whose empirical CDF $\hat{F}(z)$ should track the uniform CDF, the identity $z$, under calibration. We therefore plot the difference $\hat{F}(z) - z$ (Figure S3), together with the 95% simultaneous confidence band of Säilynoja et al. [12]. The band is calibrated by simulation so that a uniform sample stays inside it at all evaluation points $z$ jointly with probability 0.95, and it narrows to zero at the two ends because the ECDF ordinate $\hat{F}(z)$ there has vanishing binomial variance $z(1 - z)/N$. A curve contained in the band is consistent with uniformity. Finally, Table S3 reports, for each quantity, the mean and standard deviation of the ranks against their values under uniformity ($M/2 = 100$ and $\sqrt{((M + 1)^2 - 1)/12} \approx 58$), the largest absolute deviation $\max_z |\hat{F}(z) - z|$, and whether the curve stays inside the band. Because the uniformity argument assumes independent draws, we also check that thinning has removed the autocorrelation. For each quantity we estimate the effective sample size (ESS) [6] of the thinned draws and compare it to $M$, so that a ratio near one indicates the retained draws behave as independent samples. The last column of Table S3 reports the median ratio across simulations.

| Quantity | Mean rank | SD | $\max\lvert\Delta\mathrm{ECDF}\rvert$ | In band | ESS/$M$ |
|---|---|---|---|---|---|
| $\gamma$ | 99.2 | 57.9 | 0.045 | yes | 0.99 |
| $\sigma_p$ | 101.6 | 55.9 | 0.044 | yes | 1.00 |
| $\sigma_a$ | 101.2 | 58.2 | 0.046 | yes | 1.00 |
| $\sigma_\pi$ | 103.6 | 60.6 | 0.077 | yes | 1.00 |
| $p_1$ | 103.6 | 59.1 | 0.059 | yes | 1.00 |
| $a_1$ | 95.4 | 57.7 | 0.059 | yes | 0.98 |
| $\pi_{11}$ | 103.3 | 60.2 | 0.058 | yes | 1.00 |
| $\log\mu_{11}$ | 98.9 | 58.1 | 0.041 | yes | 1.00 |
| $\log\mu_{5,10}$ | 95.8 | 58.3 | 0.064 | yes | 1.00 |
| $\log\mu_{10,20}$ | 100.1 | 58.5 | 0.041 | yes | 1.00 |

TABLE S3. SBC summary over $N = 200$ simulations with $M = 200$ thinned draws. Under uniformity the rank mean is $M/2 = 100$ and the rank standard deviation is $\approx 58.0$; the Monte Carlo standard error of the mean is $\approx 4.1$. "In band" records whether the ECDF-difference curve stays within the $95\%$ simultaneous band of Säilynoja et al. [12]. $\mathrm{ESS}/M$ is the median effective-sample-size ratio of the thinned draws across simulations, where values near one confirm the draws are effectively independent. None of the quantities considered shows a departure from calibration.

#### *Results*

All 200 simulations completed and none produced divergent transitions, so the sampler traversed the posterior geometry without the failures that divergences would signal. The rank means lie between $95.4$ and $103.6$, all within one Monte Carlo standard error $(58.0/\sqrt{200} \approx 4.1)$ of the expected $100$, giving no evidence of location bias. The rank standard deviations cluster around the theoretical $58.0$, so neither over- nor under-dispersion is apparent. Every ECDF curve stays inside the simultaneous band, with the largest deviation $(0.077$, for $\sigma_\pi)$ falling near the middle of the unit interval where the band is widest. The thinning check is likewise clean, with a median ESS of at least $0.98\,M$ for every quantity, so the independence assumption behind the uniformity test holds. Figures S2 and S3 show the same picture: flat histograms and ECDF curves that fluctuates around zero without leaving the band.

For the monitored quantities the sampler reproduces the posterior implied by the model and the rank statistics show no systematic departure from uniformity, confirming that our inference is computationally faithful. As noted above, SBC assesses the correctness of the computational procedure rather than the model's fit to observed data. We therefore turn to the extensive simulation study in the next section to evaluate the model's inferential performance.

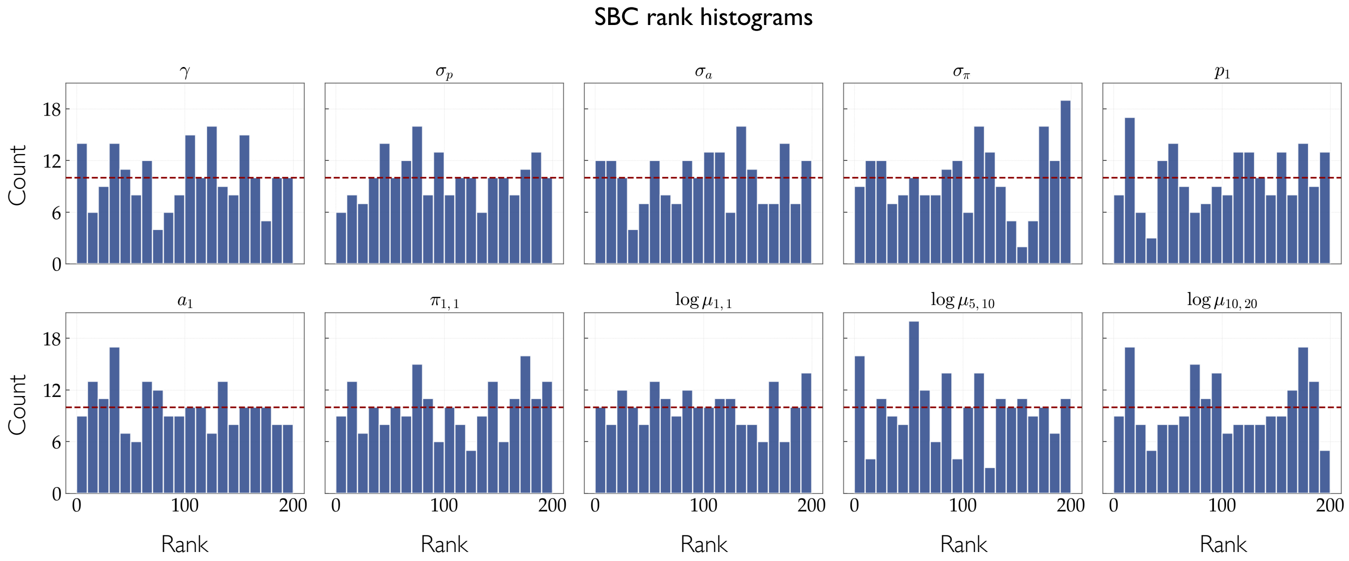


FIG. S2. **Simulation-based calibration: rank histograms.** Histograms of the simulated-parameter ranks for the ten monitored quantities, each computed from $N = 200$ simulations and aggregated into 20 equal-width bins. The dashed line marks the count expected under uniformity ($200/20 = 10$). Under calibrated inference the ranks are discrete-uniform, so an approximately flat histogram is the expected signature; systematic departures are diagnostic of specific failures, with a bowl, dome, or tilt indicating over-concentration, over-dispersion, or bias of the posterior, respectively. All ten histograms are consistent with uniformity, supporting calibrated inference across the monitored quantities.

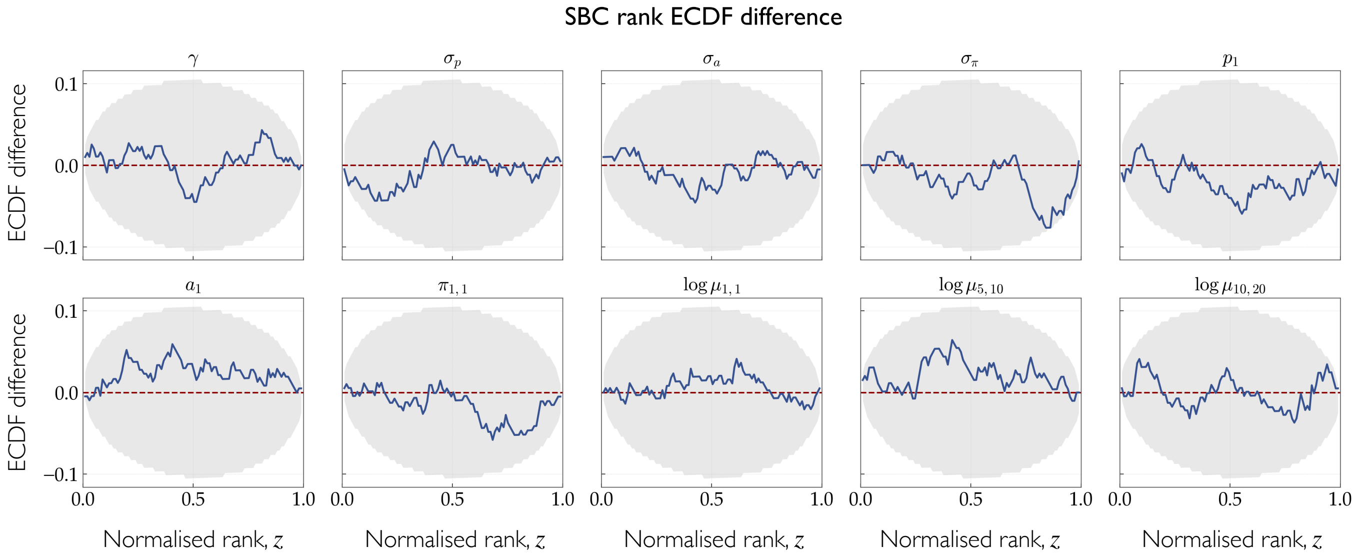


FIG. S3. **Simulation-based calibration: ECDF difference diagnostic.** For each monitored quantity, the difference $\hat{F}(z) - z$ between the empirical CDF (ECDF) of the normalized ranks $r_{n,k}/M$ and the uniform CDF expected under calibration is plot (blue curve). Subtracting the uniform CDF centres the diagnostic on zero (dotted red line, denoting perfect agreement), highlighting departures that would be hard to see in the raw ECDF. The gray region is the $95\%$ simultaneous confidence band of Säilynoja et al. [12], calibrated so that a genuinely uniform sample stays fully inside it with probability $0.95$. The band contracts to zero at the endpoints because the ECDF ordinate at $z$ has variance $z(1-z)/N$, unlike a flat Kolmogorov–Smirnov band. All curves remain within the band, consistent with uniform ranks and hence with calibrated inference.

### S7. PRIOR ROBUSTNESS

Our priors are only weakly informative, but writing them down still means fixing a few hyperparameters. In the main text we used

$$\gamma \sim N(0, \sigma_\gamma), \qquad \sigma_p, \sigma_a \sim \mathrm{Exp}(\lambda_{p,a}), \qquad \pi_{ij} \sim N(0, \sigma_\pi),$$

with $\sigma_\gamma = 3$, $\lambda_{p,a} = 2$ and $\sigma_\pi = 0.5$, where $\pi_{ij} = \log w_{ij}$ is the log-preference of plant $i$ for pollinator $j$. To check that our conclusions do not hinge on these values, we refit the model over a grid of alternatives: $\sigma_\gamma \in \{1, 2, 5\}$, $\lambda_{p,a} \in \{1, 2, 5\}$ and $\sigma_\pi \in \{0.25, 0.5, 1\}$. For each of the 27 combinations we draw 100 synthetic ground truths, fit the model to each, and measure how well we recover the quantity we actually care about, the preference matrix $w$.

Throughout the main text our point estimate for this quantity is the posterior median. For a positive, right-skewed quantity like $w$ the median is a more representative summary than the mean, which gets pulled upward by the tail, and it commutes with the log transform, so that the median preference is just the exponential of the median log-preference, and the estimate is the same whether we read it off the $w$ or the $\pi$ scale. For this reason, the first way of assess recovery we consider is the normalised mean squared error of this point estimate,

$$\mathrm{NMSE} = \frac{\sum_{ij} \big(w_{ij} - \widehat{w}_{ij}\big)^2}{\sum_{ij} w_{ij}^2},$$

with $\widehat{w}_{ij}$ the posterior median. It is 0 for perfect recovery and 1 for an estimate no better than predicting all zeros.

The second is a calibration check, closer in spirit to the Bayesian model itself. Rather than scoring a point estimate, we investigate how the posterior spread behaves in relation to the ground truth. For each entry we take the $z$-score of the ground truth against the posterior,

$$z_{ij} = \frac{\pi_{ij} - \widehat{\pi}_{ij}}{\mathrm{sd}\big(\pi_{ij}^{\mathrm{post}}\big)},$$

and average $|z_{ij}|$ over the matrix. We do this on the log scale $\pi$ rather than on $w$ since our model infers log-preferences, and the posterior is close to Gaussian, which fixes the reference value. If the posterior is well-calibrated, the ground truth behaves like a draw from it, so standardising the error by the posterior standard deviation yields $z_{ij} \sim N(0, 1)$ and hence $\mathbb{E}\,|z_{ij}| = \sqrt{2/\pi} \approx 0.8$. An average near 0.8 means the reported uncertainty matches the size of the actual error; much larger signals overconfidence, much smaller underconfidence.

The results are shown in Fig. S4, and across the whole grid the numbers stay close to each other. The NMSE stays around 0.11 (panels A–C), essentially flat over the full range of priors, so the median tracks the true preferences well regardless of the hyperparameters. The calibration diagnostic tells the same story (panels D–F), satying around its 0.8 target, drifting only gently from $\approx 0.85$ at the tightest log-preference prior ($\sigma_\pi = 0.25$) to $\approx 0.77$ at the widest ($\sigma_\pi = 1$), but never far from calibrated. The recovery of the latent preferences is therefore robust to the choice of prior hyperparameters over the range we tested.

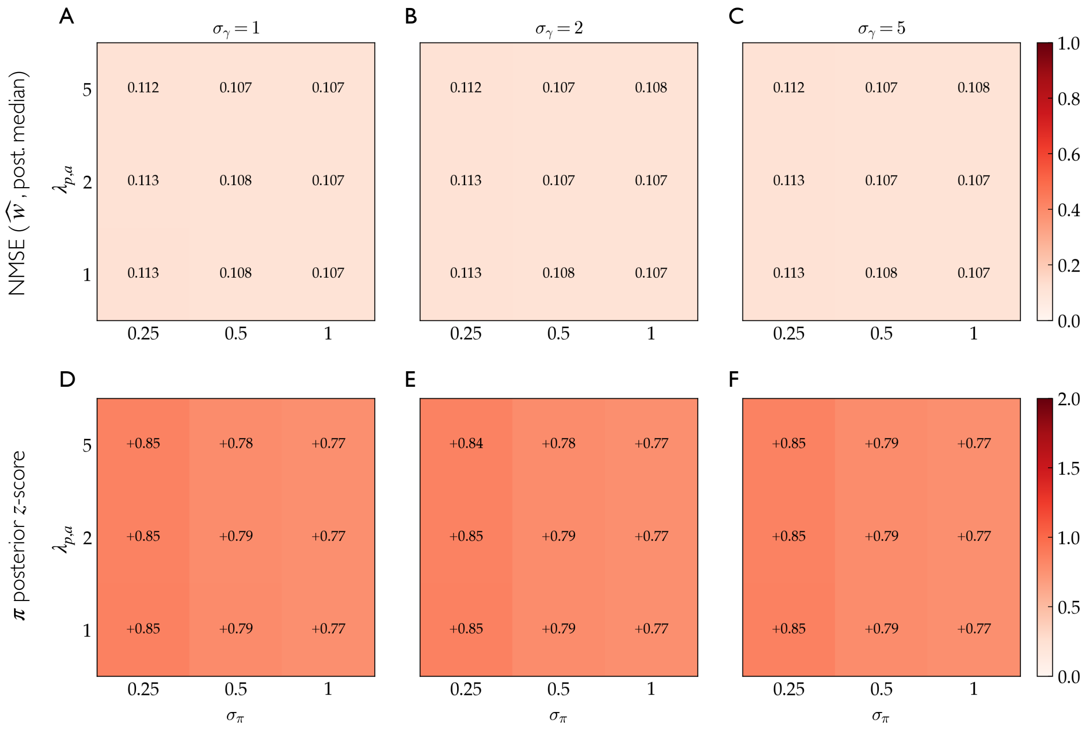


FIG. S4. **Prior robustness of the inferred preferences.** For each cell we draw 100 synthetic ground truths, refit the model, and average the metric over them. Within every heatmap the rows vary the rate of the exponential prior on the random-effect scales, $\lambda_{p,a} \in \{1, 2, 5\}$, and the columns vary the width of the log-preference prior, $\sigma_\pi \in \{0.25, 0.5, 1\}$; the three panels in each row correspond to $\sigma_\gamma \in \{1, 2, 5\}$, the width of the prior on the global intercept $\gamma$. (**A–C**) Normalised mean squared error between the posterior-median estimate and the ground-truth preferences $w$ (colour and printed value); 0 is perfect recovery. (**D–F**) Mean absolute $z$-score of the ground truth against the posterior, computed on the log-scale $\pi$; the target for a well-calibrated Gaussian posterior is $\approx 0.8$. Both metrics are near-constant across all 27 hyperparameter combinations, and close to the target values, showing good recovery and calibration, insensitive to the prior.

### S8. MODEL RECOVERY PERFORMANCE

#### A. Single-observation model

For each quantity we summarise the model recovery using the posterior *median* as point estimate $\hat{\theta}$ and the following metrics:

1. $\gamma$: relative error, $|\hat{\gamma} - \gamma|/|\gamma|$ (replaced by $|\hat{\gamma} - \gamma|$ when $\gamma = 0$);

2. $\boldsymbol{\pi}$: Hamming distance and Spearman correlation between $\widehat{\boldsymbol{\pi}}$ and $\boldsymbol{\pi}$. These two metrics assess complementary aspects of the recovered log-preferences, whose sign encodes whether a pair is preferred ($\pi_{ij} > 0$) or avoided ($\pi_{ij} < 0$). The Hamming distance is the sign-recovery error, i.e. the fraction of entries whose estimated sign disagrees with the truth,

$$\mathrm{Ham} = \frac{1}{|\mathcal{S}|} \sum_{(i,j)\in\mathcal{S}} \mathbb{I}\big[\mathrm{sign}\, \widehat{\pi}_{ij} \neq \mathrm{sign}\, \pi^{\star}_{ij}\big], \qquad \mathcal{S} = \{(i,j) : |\pi^{\star}_{ij}| \geq \delta\}, \tag{S7}$$

restricted to entries with $|\pi^{\star}_{ij}| \geq \delta$ ($\delta = 0.1$), since near-zero preferences have no well-defined direction and their sign would be set by noise. This asks only whether each meaningful pair is placed on the correct side of the baseline, ignoring magnitude and ordering, and takes values in $[0, 1]$, with 0 for perfect sign recovery and $\approx 0.5$ expected from random signs. The Spearman ($\rho$, in $[-1, 1]$) correlation instead compares the full vector of inferred and true log-preferences, specifically measuring the ordering of the pairs by preference strength. Reporting both allows to separates recovery of direction from recovery of relative strength.

3. $\boldsymbol{\sigma}$ and $\boldsymbol{\tau}$: squared Jensen–Shannon divergence. For the species-level effects we take the recovered $\boldsymbol{\sigma} = \exp(\mathbf{p})$ and $\boldsymbol{\tau} = \exp(\mathbf{a})$ and normalise them to sum to one, $\boldsymbol{\sigma}/\sum_i \sigma_i$ and $\boldsymbol{\tau}/\sum_j \tau_j$, so that each defines an abundance distribution over plants and over pollinators. The squared Jensen–Shannon divergence then measures how far the recovered distribution is from the truth. This measure is symmetric and bounded, with base-2 logarithms it lies in $[0, 1]$, equal to 0 for identical distributions and approaching 1 only when their supports are essentially disjoint. Working on the normalised scale makes the recovery interpretable as recovery of the abundance *shape*, i.e. which species are common and which are rare. We do not apply it to $\mathbf{p}$ or $\mathbf{a}$ directly, as those are zero-centred log-effects rather than distributions.

4. Additionally we consider NRMSE (see below) for the composite variable $\log \mu$.

For $\gamma, \mathbf{p}, \mathbf{a}$ and $\boldsymbol{\pi}$ we also compute the credible-interval coverage, the relative bias, and the (normalised) root mean squared error. For an array-valued parameter $\theta = (\theta_1, \ldots, \theta_n)$ with posterior median estimate $\hat{\theta}$ and ground truth $\theta^{\star}$, these are defined as follows.

The coverage is the fraction of components whose true value lies inside the elementwise 95% credible interval $[q_i^{2.5}, q_i^{97.5}]$,

$$\mathrm{cov}(\theta) = \frac{1}{n} \sum_{i=1}^{n} \mathbb{I}\big[\, q_i^{2.5} \leq \theta^{\star}_i \leq q_i^{97.5} \,\big], \tag{S8}$$

which under correct calibration is expected to equal 0.95. The relative bias normalises the *net* signed error by the summed absolute magnitude of the ground truth [9],

$$\mathrm{relbias}(\theta) = \frac{\sum_{i=1}^{n} \big(\hat{\theta}_i - \theta^{\star}_i\big)}{\sum_{i=1}^{n} \big|\theta^{\star}_i\big|}, \tag{S9}$$

a stable choice for the zero-centred parameters $p, a, \pi$, for which an elementwise ratio would be ill-defined. The root mean squared error and its normalised counterpart are

$$\mathrm{RMSE}(\theta) = \sqrt{\frac{1}{n} \sum_{i=1}^{n} \big(\hat{\theta}_i - \theta^{\star}_i\big)^2}, \qquad \mathrm{NRMSE}(\theta) = \frac{\mathrm{RMSE}(\theta)}{\max(\theta^{\star}) - \min(\theta^{\star})}, \tag{S10}$$

where we normalise for the range of ground truth values, so that NRMSE expresses the error in units of the spread of $\theta^{\star}$ and is comparable across parameters.

Specifically, we conduct the following analyses:

1. we vary the heterogeneity of the species abundance distribution, $\sigma_{\text{abund}}$ (which sets $\sigma_p$ and $\sigma_a$, so that $p, a \sim \mathcal{N}(0, \sigma^2_{\text{abund}})$), over the values $\{0.1, 0.5, 1, 2\}$;
2. we vary the pairwise (log-)preference variance, $\sigma_\pi \in \{0.1, 0.5, 1, 1.5\}$, which controls the heterogeneity of the preferences (and, at low values, also their magnitude);
3. we repeat both analyses for two network sizes, $10 \times 20$ and $20 \times 60$, chosen to mirror the plant-to-pollinator ratio typically observed in empirical data.

*Results*

The four sweeps in Fig. S5 confirm that recovery is accurate across most of the parameter space and isolate where, and why, it deteriorates. The dominant axis is the baseline $\gamma$, which controls the grand total of observed counts (leftmost column, growing by orders of magnitude with $\gamma$). More interactions mean more information, and every metric improves monotonically with $\gamma$. The clearest instance is the sign of the pairwise preferences: the Hamming distance on $\boldsymbol{\pi}$ is high at the lowest baseline but collapses to near zero once $\gamma$ crosses a moderate value, above which signs are recovered almost perfectly, while the Spearman correlation on $\boldsymbol{\pi}$ rises in parallel, indicating that both the direction and the relative ordering of preferences become well identified. The baseline $\gamma$ itself is recovered with small relative error throughout, except at $\gamma = -1$; notably, its recovery is degraded by species-effect heterogeneity, the relative error on $\gamma$ grows with $\sigma_{\text{abund}}$ (top two rows), but is essentially insensitive to preference heterogeneity (bottom two rows). Species activity distributions $\boldsymbol{\sigma}$ and $\boldsymbol{\tau}$ are recovered well in all conditions, their Jensen–Shannon divergence satying always close to zero. The NRMSE on $\log \mu$ likewise decreases steadily towards zero as $\gamma$ increases, and it is never higher than $0.2$. Preference heterogeneity has little effect on its own, with one exception: at $\sigma_\pi = 0.1$ the preferences are so narrow and weak that the signal is close to absent, which lowers preference recovery (higher Hamming, lower Spearman) at low baseline—the model cannot recover structure that is barely present in the ground truth. Contrasting the two heterogeneity sweeps, variability in species activity and variability in preferences act on different targets: the former mainly perturbs the baseline, whereas the latter matters only when preferences are too small to be detected. Finally, comparing the $10 \times 20$ and $20 \times 60$ sweeps, the larger networks yield a modest, uniform improvement—slightly lower errors and smaller variability across realisations—but no qualitative change, indicating that the conclusions are not an artefact of a particular network size. In short, the model correctly recovers the baseline, species-level effects, and pairwise preferences almost everywhere, with degradation confined to the expected data-poor or weak-signal extremes ($\gamma = -1$, or $\sigma_\pi = 0.1$ at low baseline).

Figures S6–S9 report the coverage, relative bias and (n)RMSE of $\gamma$, $\mathbf{p}$, $\mathbf{a}$ and $\boldsymbol{\pi}$, for varying abundances (network sizes $10 \times 20$ and $20 \times 60$) and varying preferences (both sizes again), as a function of the true baseline $\gamma$ ($x$-axis) and of the swept heterogeneity parameter ($y$-axis). When varying abundances, for both network sizes we obtain coverage close to the nominal $95\%$, small relative bias and near-zero NRMSE, with two exceptions: the RMSE on $\gamma$ grows towards the high-baseline end ($\gamma = 5$), and the coverage and bias of the individual species effects ($p$, $a$) deteriorate mildly at the highest species-effect heterogeneity ($\sigma_{\text{abund}} = 2$). Our main quantity of interest, the pairwise preferences $\boldsymbol{\pi}$, is recovered with nominal coverage, zero bias and very low error throughout the abundance sweep. When varying preferences, and consistently with the earlier analyses, the swept parameter has a weaker effect on these metrics. The relative bias is essentially uniform across parameter space and shows no pathological behaviour, and the NRMSE remains low, decreasing with the baseline as expected. Coverage degrades slightly for the larger network ($20 \times 60$), though it never drops below roughly $90\%$, whereas for $10 \times 20$ it is generally closer to nominal for $\gamma$, $p$ and $a$ but worse for $\boldsymbol{\pi}$ at the lowest preference heterogeneity ($\sigma_\pi = 0.1$), where the preference signal is weakest.

#### B. Multi-site model

We repeat the recovery analysis of the previous section for the multi-site model, using the same point estimate (posterior median) and the same metrics for $\gamma$, $\bar{\mathbf{p}}$, $\bar{\mathbf{a}}$ and $\boldsymbol{\pi}$. We additionally assess the recovery of the site-level deviations of the core species effects: $\delta^p$ (an $n_{\text{plants}} \times n_{\text{sites}}$ matrix giving, for each site, the departure of the plant effects from their core value $\bar{\mathbf{p}}$) and $\delta^a$ (the $n_{\text{polls}} \times n_{\text{sites}}$ analogue for pollinators). For each we report the Spearman correlation between inferred and true values, assessing how well the per-site structure is ordered, in addition to the coverage, relative bias and (N)RMSE computed for all array parameters.

Because the single-site sweep showed only a modest, qualitatively neutral effect of network size (Fig. S5), and to keep the already large multi-site grid tractable, we fix the network to a single size, $10 \times 20$, and sweep the baseline $\gamma$, the species-effect heterogeneity $\sigma_{\text{abund}}$, and the number of sites $n_{\text{sites}} \in \{3, 5, 10\}$; the single-site fit is overlaid as a reference. We keep the abundance sweep rather than the preference sweep, as the former was the one modulating recovery in the single-site analysis, whereas preference heterogeneity had little effect except in the degenerate weak-signal case.

*Results*

We obtain a similar picture to that of the single-site analysis (Fig. S10). Recovery again improves monotonically with the baseline $\gamma$: the Hamming distance on $\boldsymbol{\pi}$ collapses and the Spearman correlation on $\boldsymbol{\pi}$ rises as $\gamma$ grows, while the Jensen–Shannon divergences on $\boldsymbol{\sigma}$ and $\boldsymbol{\tau}$ stay near zero and the NRMSE on $\log \mu$ decreases steadily throughout. Note that here the grand total in the leftmost column is summed over entries and over sites, $\sum_s \sum_{ij} M_{ij}^{(s)}$, so it reaches appreciably higher values than in the single-site case. This is most pronounced at $\gamma = 5$, an extreme baseline we include to stress-test the model but that represents a very high sampling intensity and community baseline, unlikely to be reached in practice. The site-level deviations $\delta^p$ and $\delta^a$ are recovered well across conditions, their Spearman correlation is high over most of the grid and is largely insensitive to the number of sites, so that three sites already suffice and moving to ten changes little. As in the single-site case, the effect of species-effect heterogeneity is concentrated on the baseline and, at low counts, on the species distributions, while the preference recovery is governed mainly by $\gamma$. The main multi-site–specific signal is the interaction between sites and baseline: at low per-site counts, adding sites appreciably improves recovery of the pairwise preferences $\boldsymbol{\pi}$ relative to the single-site reference, whereas once the baseline is high a single well-sampled site does about as well and extra sites add little; this gain from additional sites is larger at higher $\sigma_{\mathrm{abund}}$. Overall, the multi-site model recovers the baseline, core species effects, site deviations and pairwise preferences accurately across the grid, with degradation again confined to the data-poor extreme (low $\gamma$).

Pooling over $\sigma_{\mathrm{abund}}$ (Fig. S11), coverage sits close to the nominal $95\%$, relative bias stays small and the NRMSE is low for every parameter, with little dependence on the number of sites. The only appreciable degradation appears at the extreme baseline $\gamma = 5$—in the recovery of $\gamma$ itself (lower coverage and higher RMSE) and, more mildly, in the coverage of the core species effects—which nonetheless corresponds to a very high sampling intensity unlikely to be reached in practice. This suggests that at such high counts the model struggles slightly to separate what is due to the overall baseline from what is due to species individual activity levels.

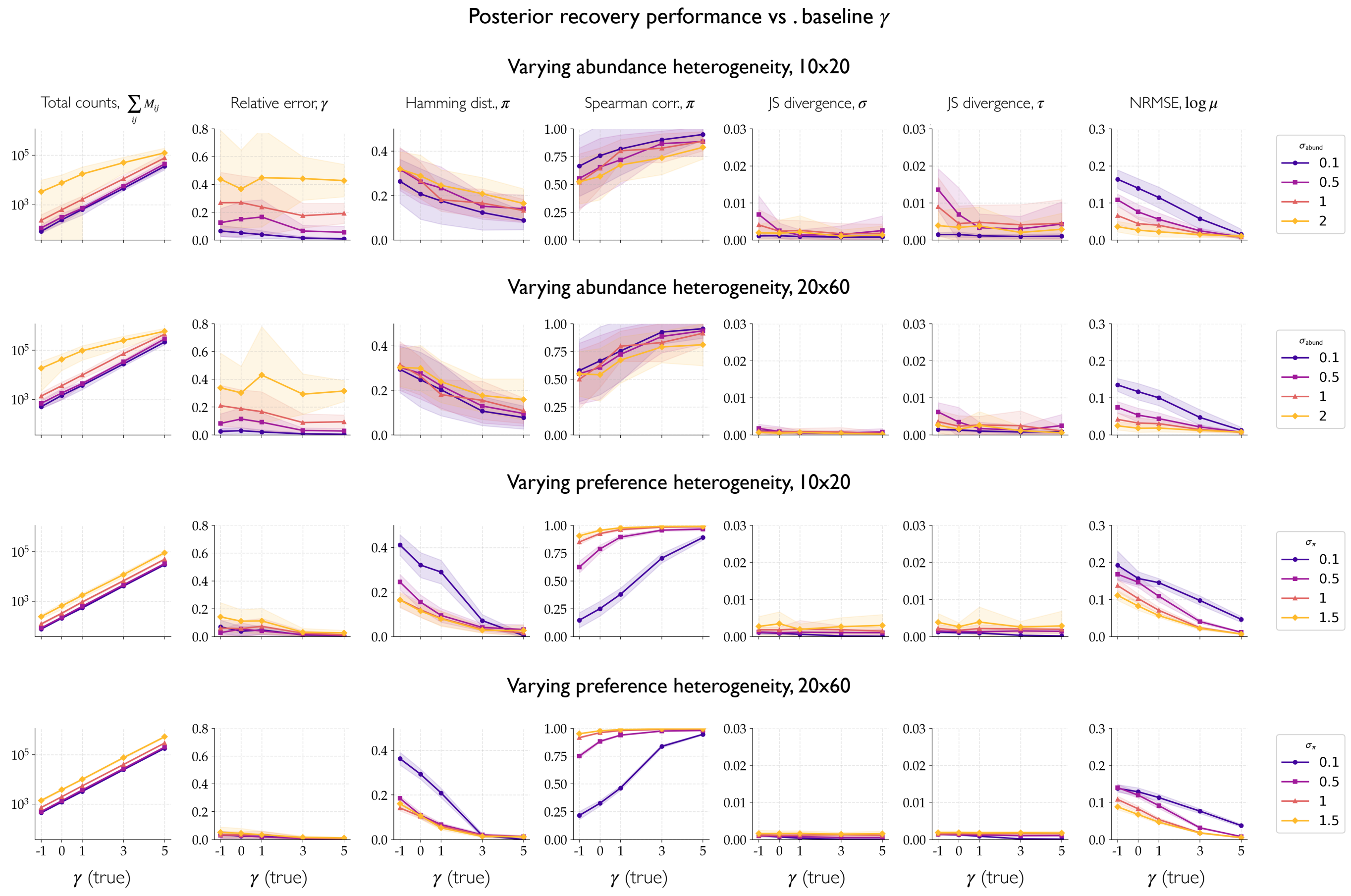


FIG. S5. **Recovery performance across parameter space.** Model performance in recovering ground-truth parameters from synthetic data, as a function of the true baseline $\gamma$ ($x$-axis of every panel). Rows correspond to the four sweeps: varying species-effect heterogeneity $\sigma_{\text{abund}}$ at network size $10 \times 20$ and $20 \times 60$ (top two rows), and varying preference heterogeneity $\sigma_\pi$ at the same two sizes (bottom two rows); colour encodes the swept parameter ($\sigma_{\text{abund}}$ or $\sigma_\pi$, see legends). For each parameter configuration, 50 independent synthetic datasets are generated, with stochasticity arising from the data-generating process. For each dataset the model is fit 20 times and the point estimate is taken as the median of the 20 posterior medians, to average out sampling variability across runs. Each point is the resulting accuracy measure averaged over the 50 realisations, and shaded regions show the standard deviation across realisations. The leftmost column reports the grand total of the simulated count matrices, $\sum_{ij} M_{ij}$ (mean and standard deviation), for each baseline value; the remaining columns, from left to right, report the relative error on $\gamma$, the Hamming distance and Spearman correlation on $\boldsymbol{\pi}$, the Jensen–Shannon divergence on $\boldsymbol{\sigma}$ and $\boldsymbol{\tau}$, and the NRMSE on $\log \mu$. Across the whole grid the model recovers all quantities accurately, with small errors for most configurations. Performance degrades appreciably only in data-poor or weak-signal corners, very low baseline ($\gamma = -1$), strong species-effect heterogeneity, or near-flat preferences ($\sigma_\pi = 0.1$), and improves smoothly as the baseline, and hence the total number of observed interactions, grows.

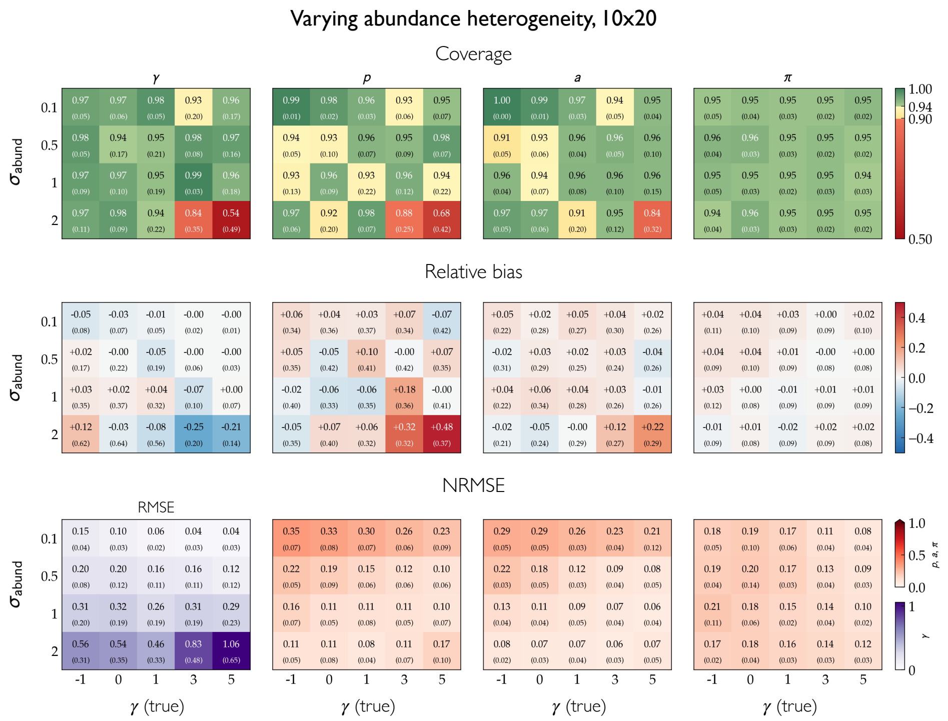


FIG. S6. **Recovery diagnostics for the abundance sweep,** $10 \times 20$ **networks.** Coverage of the central 95% credible interval (top), relative bias (middle) and RMSE/NRMSE (bottom) for the baseline $\gamma$, the species effects $\mathbf{p}$ and $\mathbf{a}$, and the pairwise preferences $\boldsymbol{\pi}$, as a function of the true baseline $\gamma$ ($x$-axis) and the species-effect heterogeneity $\sigma_{\text{abund}}$ ($y$-axis). Each cell averages over the 50 realisations; the value below each entry is the standard deviation across realisations. Coverage panels share a colour scale centred on the nominal 0.95; NRMSE is shown for $\mathbf{p}$, $\mathbf{a}$, $\boldsymbol{\pi}$ and absolute RMSE (purple scale) for $\gamma$.

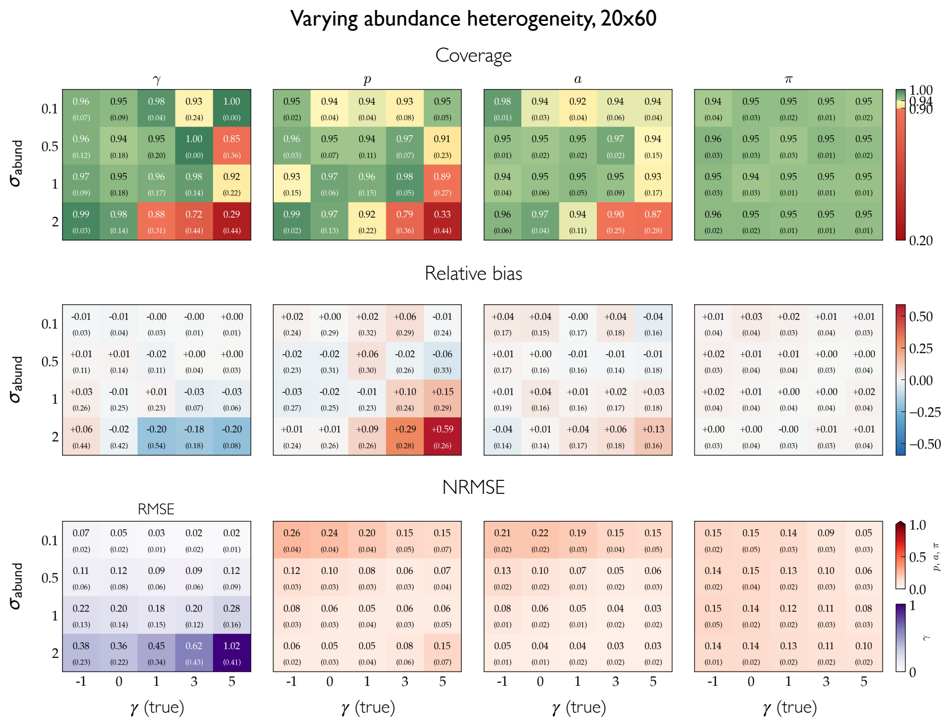


FIG. S7. **Recovery diagnostics for the abundance sweep,** $20 \times 60$ **networks.** As in Fig. S6, for the larger network size. Coverage (top), relative bias (middle) and RMSE/NRMSE (bottom) for $\gamma$, $\mathbf{p}$, $\mathbf{a}$ and $\boldsymbol{\pi}$, against the true baseline $\gamma$ ($x$-axis) and the species-effect heterogeneity $\sigma_{\text{abund}}$ ($y$-axis); cell values are means over the 50 realisations, with the standard deviation below.

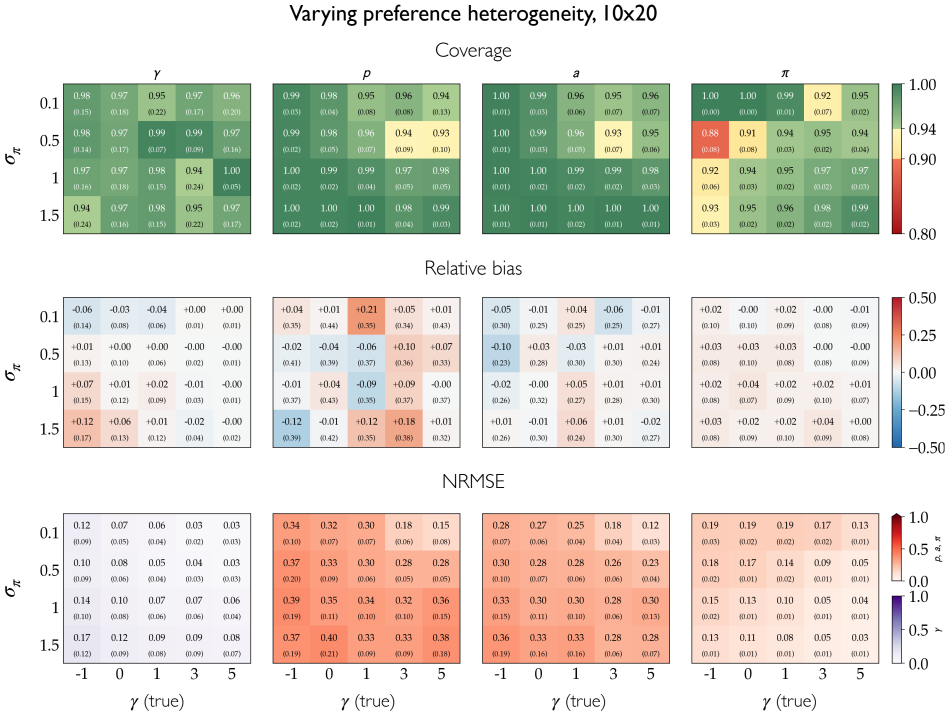


FIG. S8. **Recovery diagnostics for the preference sweep,** $10 \times 20$ **networks.** As in Fig. S6, but sweeping the preference heterogeneity $\sigma_\pi$ ($y$-axis) instead of $\sigma_{\mathrm{abund}}$. Coverage (top), relative bias (middle) and RMSE/NRMSE (bottom) for $\gamma$, $\mathbf{p}$, $\mathbf{a}$ and $\boldsymbol{\pi}$, against the true baseline $\gamma$ ($x$-axis); cell values are means over the 50 realisations, with the standard deviation below.

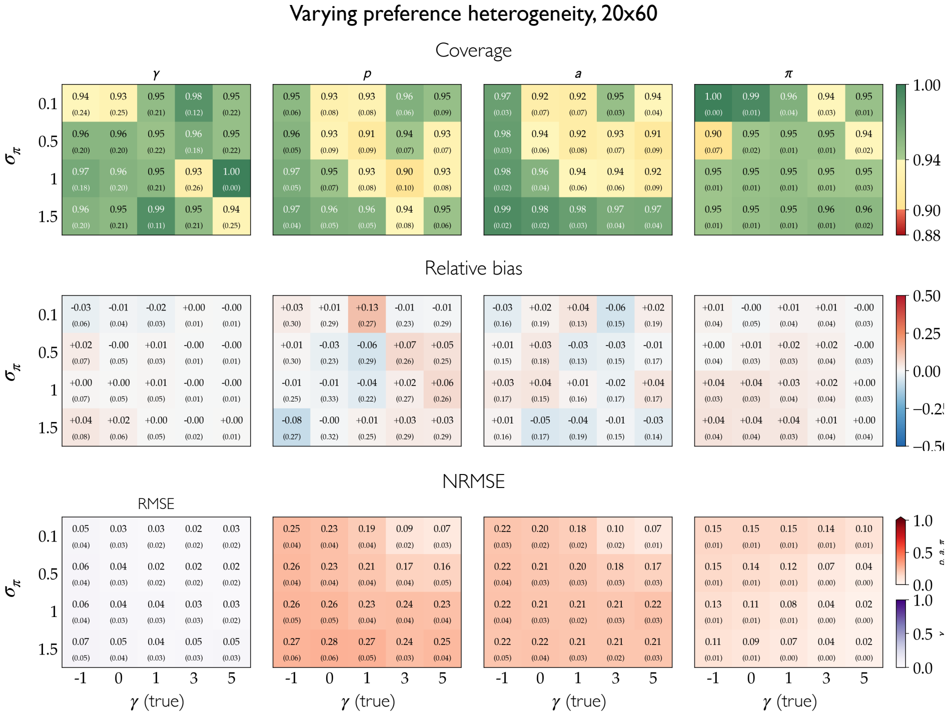


FIG. S9. **Recovery diagnostics for the preference sweep,** $20 \times 60$ **networks.** As in Fig. S8, for the larger network size. Coverage (top), relative bias (middle) and RMSE/NRMSE (bottom) for $\gamma$, $\mathbf{p}$, $\mathbf{a}$ and $\boldsymbol{\pi}$, against the true baseline $\gamma$ ($x$-axis) and the preference heterogeneity $\sigma_\pi$ ($y$-axis); cell values are means over the 50 realisations, with the standard deviation below.

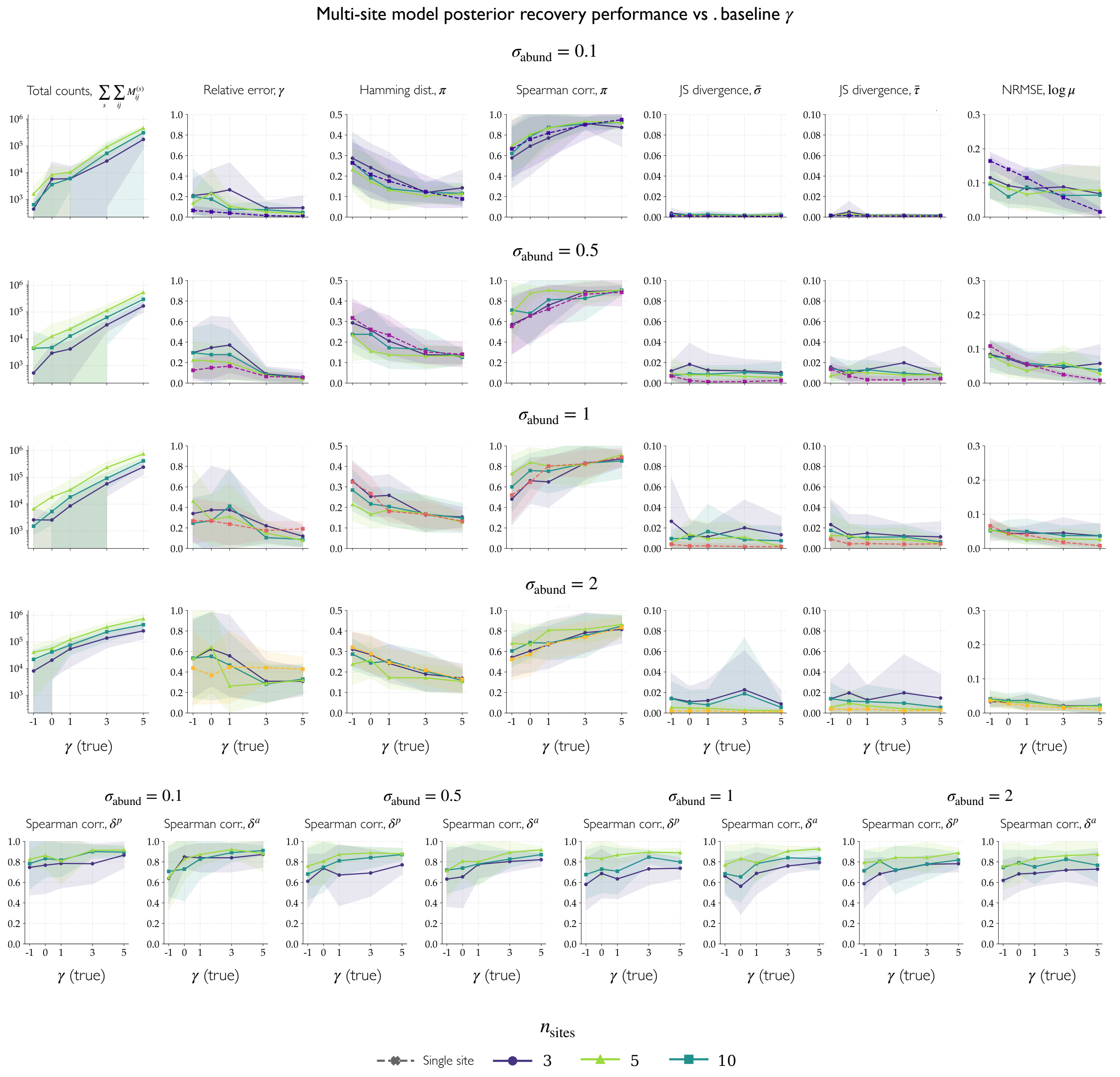


FIG. S10. **Recovery performance of the multi-site model.** Recovery of the ground-truth parameters as a function of the true baseline $\gamma$ ($x$-axis of every panel), for $10 \times 20$ network size. Rows correspond to the swept species-effect heterogeneity $\sigma_{\text{abund}} \in \{0.1, 0.5, 1, 2\}$; coloured lines give the number of sites $n_{\text{sites}}$ with the single-site fit overlaid as a dashed reference (see legend). For each cell, 30 ground truths are drawn and, for each, 10 count matrices are simulated and fit; each point is the metric averaged first over replicates and then over ground truths, and shaded regions show the standard deviation across ground truths. The leftmost column reports the grand total of simulated counts across sites $\sum_s \sum_{ij} M_{ij}^{(s)}$; the remaining columns of the upper block report, from left to right, the relative error on $\gamma$, the Hamming distance and Spearman correlation on $\boldsymbol{\pi}$, the Jensen–Shannon divergence on $\bar{\boldsymbol{\sigma}}$ and on $\bar{\boldsymbol{\tau}}$, and the NRMSE on $\log \mu$. The bottom row shows the Spearman correlation on the site-level deviations $\delta^p$ and $\delta^a$, one pair per value of $\sigma_{\text{abund}}$ (indicated above each pair), where $\delta^p$ and $\delta^a$ are the per-site departures of the plant and pollinator effects from their core values.

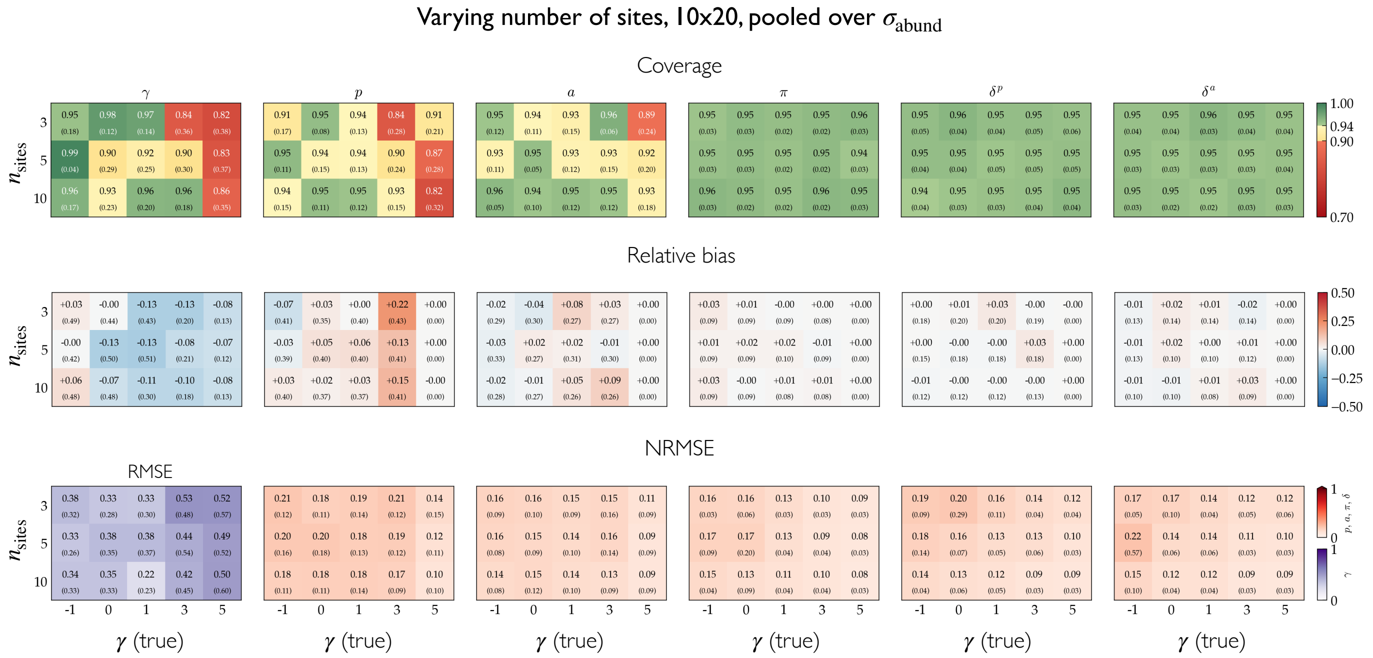


FIG. S11. **Recovery diagnostics of the multi-site model, pooled over species-effect heterogeneity.** Coverage (top), relative bias (middle) and NRMSE (bottom) for the baseline $\gamma$, the core species effects $\bar{\mathbf{p}}$ and $\bar{\mathbf{a}}$, the pairwise preferences $\boldsymbol{\pi}$, and the site-level deviations $\delta^p$ and $\delta^a$, as a function of the true baseline $\gamma$ ($x$-axis) and the number of sites $n_{\mathrm{sites}}$ ($y$-axis). Each cell pools all ground truths across the four $\sigma_{\mathrm{abund}}$ values: the value is the mean over the pooled ground truths and the number below is the corresponding standard deviation. $\gamma$ is scored by RMSE (purple scale) rather than nRMSE, as it is a single latent-scale parameter. Recovery is accurate across the grid, coverage close to the nominal 95%, small bias and low error, and is largely insensitive to $n_{\mathrm{sites}}$.

### S9. SINGLE-OBSERVATION EMPIRICAL CASE STUDY

For the single-observation empirical case study (Fig. 4), all convergence diagnostics were satisfactory (Table S4). The rank-normalized split-$\hat{R}$ was $1.000$ for every parameter, and both bulk- and tail-effective sample sizes were well in excess of the target of $400$ across all parameter blocks (minimum bulk-ESS $4283$, minimum tail-ESS $4381$). No transition was divergent or saturated the maximum tree depth, and the energy diagnostic (E-BFMI) was satisfactory throughout.

| Parameter | $\widehat{R}$ | bulk-ESS | tail-ESS |
|---|---:|---:|---:|
| $\gamma$ | 1.000 | 2403 | 2516 |
| $p_i$ | 1.000 | 2139–4088 | 2697–3547 |
| $a_j$ | 1.000 | 1151–4657 | 927–3459 |
| $\pi_{ij}$ | 1.000–1.010 | 1118–8984 | 1429–3419 |
| $\sigma_p,\ \sigma_a,\ \sigma_\pi$ | 1.000 | 997–2104 | 808–2933 |

TABLE S4. Convergence diagnostics for the single-observation empirical case study, reported by parameter block: the baseline $\gamma$, plant effects $p_i$, pollinator effects $a_j$, pairwise preferences $\pi_{ij}$, and the random-effect scales $\sigma_p, \sigma_a, \sigma_\pi$. Values are ranges (minimum–maximum) across the parameters in each block, except for the scalar $\gamma$. All $\widehat{R} = 1.000$ and all effective sample sizes exceed 400.

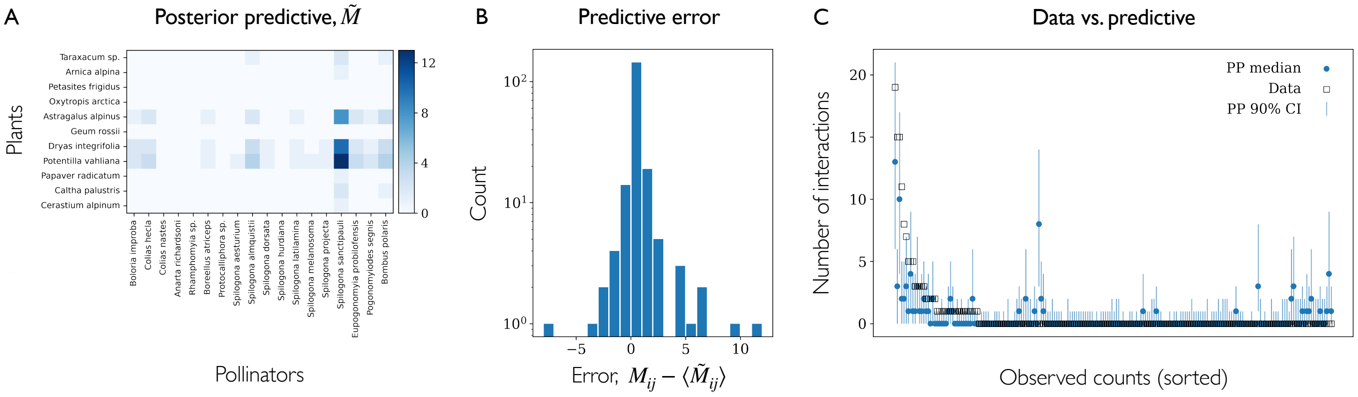


FIG. S12. **Posterior predictive check for the reduced model without pair-specific preferences.** We refit the model to the empirical data used in the main text (dataset ID "MOMA" from [11]) removing the pair-specific preference term $\pi_{ij}$, and repeat the posterior predictive check. (**A**) Posterior-predictive median counts $\widetilde{M}_{ij}$ for every plant-pollinator pair. The lack of a pair-specific interaction term is already visible in the pattern of this matrix, which is dominated by the rows and columns corresponding to the more active species. (**B**) Distribution of the predictive error $M_{ij} - \langle \widetilde{M}_{ij} \rangle$ between observed counts and their posterior-predictive values, pooled over all pairs (note the logarithmic count axis). (**C**) Observed counts (open squares) against the posterior-predictive median (filled circles) and 90% credible interval (vertical bars) for each pair, ordered by decreasing observed count. Compared with the full model in the main text, the reduced model reproduces the data markedly less well. Errors are more widely spread and many observed counts, particularly the most frequent interactions, fall outside their 90% predictive intervals, confirming that the pair-specific preference term is needed.

### S10. MULTI-OBSERVATION EMPIRICAL CASE STUDY

For the multi-observation case study, we ran 2000 post-warmup iterations per chain (4 chains) to accommodate the autoregressive temporal component, whose correlation and variance parameters mix more slowly. Convergence diagnostics are reported in Table S5. The rank-normalized split-$\widehat{R}$ did not exceed 1.01 for any parameter, and tail effective sample sizes were above 400 throughout. Bulk effective sample sizes exceeded 400 for all parameters except the plant-effect scale $\sigma_p$ (341) and the temporal-correlation parameter $\rho$ (364), both slightly below the target; the parameters of primary interest—the pairwise preferences $\pi_{ij}$ and the species- and site-level effects—were well sampled, with bulk-ESS in the thousands. The energy diagnostic (E-BFMI) was satisfactory, no transition saturated the maximum tree depth, and 3 of 2000 transitions (0.15%) were divergent, a negligible fraction that does not affect the reported summaries.

| Parameter | $\widehat{R}$ | bulk-ESS | tail-ESS |
|---|---:|---:|---:|
| $\bar{\gamma}$ | 1.000 | 1842 | 1239 |
| $\delta_s^\gamma$ | 1.000 | 1873–2585 | 1150–1209 |
| $\eta_t^\gamma$ | 1.000 | 2097–6840 | 3129–4878 |
| $\bar{p}_i$ | 1.000–1.010 | 546–7740 | 3271–5452 |
| $\bar{a}_j$ | 1.000 | 719–11698 | 1087–6908 |
| $\delta_{i,s}^p$ | 1.000 | 2770–11305 | 3520–6840 |
| $\delta_{j,s}^a$ | 1.000 | 4299–22403 | 4672–7550 |
| $\eta_{i,t}^p$ | 1.000–1.010 | 505–10253 | 1487–6628 |
| $\eta_{j,t}^a$ | 1.000 | 607–19834 | 787–7370 |
| $\pi_{ij}$ | 1.000–1.010 | 1555–25308 | 3145–7402 |
| $\sigma_p$ | 1.010 | **341** | 1469 |
| $\sigma_a$ | 1.000 | 730 | 983 |
| $\sigma_\pi$ | 1.000 | 3768 | 5612 |
| $\Sigma_\gamma$ | 1.000 | 1962 | 1261 |
| $\Sigma_p$ | 1.000 | 4093 | 4833 |
| $\Sigma_a$ | 1.000 | 3307 | 5720 |
| $\Sigma_{\gamma,t}$ | 1.000 | 1315 | 3688 |
| $\Sigma_{p,t}$ | 1.010 | 516 | 1159 |
| $\Sigma_{a,t}$ | 1.010 | 462 | 973 |
| $\rho$ | 1.010 | **364** | 772 |

TABLE S5. Convergence diagnostics for the multi-observation case study, reported by parameter block. Values are ranges (minimum–maximum) across the parameters in each block, except for scalar parameters. Blocks comprise the intensity terms ($\bar{\gamma}$ and its site and temporal deviations $\delta_s^\gamma, \eta_t^\gamma$), the core species effects ($\bar{p}_i, \bar{a}_j$) with their site and temporal deviations ($\delta_{i,s}^p, \delta_{j,s}^a, \eta_{i,t}^p, \eta_{j,t}^a$), the pairwise preferences $\pi_{ij}$, the random-effect scales ($\sigma_p, \sigma_a, \sigma_\pi$ and the deviation scales $\Sigma_p, \Sigma_a, \Sigma_{p,t}, \Sigma_{a,t}$), and the temporal-correlation parameter $\rho$. The two bulk-ESS values below 400 (bold) correspond to variance/correlation parameters, all $\widehat{R} \leq 1.01$.

### S11. URBANISATION GRADIENT CASE STUDY

#### A. Full model description

Here we describe the specific parametrization of the model used for the urbanisation case study, which lets a single binary covariate act on *every* level of the model, in the spirit of a two-group treatment contrast. The urbanisation class is recorded as an ordinal variable (low, medium, high). To obtain a clean two-group contrast we retain only the extreme classes and define the binary site-level covariate $x_s = 1$ for high-urbanisation sites and $x_s = 0$ for low-urbanisation sites, discarding the intermediate sites. The baselines therefore describe the low-urbanisation group, and every urbanisation parameter is a low$\rightarrow$high contrast on the log scale.

The log interaction rate becomes

$$\begin{aligned}\log \mu_{ij}^{(s,t)} = &\underbrace{\bar{\gamma} + \delta_s^{\gamma} + \beta^{\gamma}\, x_s}_{\text{baseline}} \\ &+ \underbrace{\bar{p}_i + \eta_{i,t}^{p} + \beta_i^{p}\, x_s}_{\text{plant activity}} \\ &+ \underbrace{\bar{a}_j + \eta_{j,t}^{a} + \beta_j^{a}\, x_s}_{\text{pollinator activity}} \\ &+ \underbrace{\pi_{ij} + \beta_{ij}^{\pi}\, x_s}_{\text{preference}}, \end{aligned} \tag{S11}$$

where the terms $\bar{\gamma}, \delta_s^{\gamma}, \bar{p}_i, \eta_{i,t}^{p}, \bar{a}_j, \eta_{j,t}^{a}, \pi_{ij}$ retain the hierarchical priors and AR(1) temporal structure of the full model as described in the main text. For completeness we report the priors on these terms. The global intercept is $\bar{\gamma} \sim \mathcal{N}(0, 2^2)$ and the site deviations are $\delta_s^{\gamma} \sim \mathcal{N}(0, \Sigma_{\gamma}^2)$ with $\Sigma_{\gamma} \sim \text{Exp}(2)$. The species cores are $\bar{p}_i \sim \mathcal{N}(0, \sigma_p^2)$ and $\bar{a}_j \sim \mathcal{N}(0, \sigma_a^2)$, with $\sigma_p, \sigma_a \sim \text{Exp}(2)$. The temporal deviations $\eta_{i,t}^{p}$ and $\eta_{j,t}^{a}$ follow the stationary AR(1) process of Eqs. (10), with shared autocorrelation $\rho \sim \mathcal{N}(0.5, 0.3^2)$ truncated to $(-1, 1)$ and $\Sigma_{p,t}, \Sigma_{a,t} \sim \text{Exp}(2)$. The latent preferences are $\pi_{ij} \sim \mathcal{N}(0, \sigma_{\pi}^2)$ with the half-normal hyperprior $\sigma_{\pi} \sim \mathcal{N}^{+}(0, 0.5^2)$.

Because the design collapses to two urbanisation groups, we do not fit free per-site species deviations $\delta_{i,s}^{p}$; the species-level between-site structure is instead carried entirely by the urbanisation contrast, which is both more parsimonious given the small number of sites and directly interpretable as the quantity of interest.

The four urbanisation channels are ecologically distinct. The scalar $\beta^{\gamma}$ is an unconstrained shift capturing the average change in overall interaction intensity between the two groups. The vectors $\beta_i^{p}$ and $\beta_j^{a}$ capture differential, species-specific changes in activity (effective abundance) under urbanisation. The matrix $\beta_{ij}^{\pi}$ captures *rewiring*, that is, changes in pair-specific preference that are not attributable to shifts in species activity.

Since the channels are additive on the log scale, they are not jointly identified without constraints. Any constant added to $\beta^{p}$ (or $\beta^{a}$) could be reabsorbed into $\beta^{\gamma}$, so we impose sum-to-zero constraints, $\sum_i \beta_i^{p} = 0$ and $\sum_j \beta_j^{a} = 0$, forcing these channels to carry only differential (mean-removed) abundance changes. Likewise, the row and column means of $\beta^{\pi}$ would be confounded with the two activity contrasts, so we double-centre the rewiring block, imposing $\sum_i \beta_{ij}^{\pi} = 0$ and $\sum_j \beta_{ij}^{\pi} = 0$. Concretely, an unconstrained matrix $B$ is centred as

$$B_{ij}^{*} = B_{ij} - \bar{B}_{i\bullet} - \bar{B}_{\bullet j} + \bar{B}_{\bullet\bullet}, \tag{S12}$$

with $\bar{B}_{i\bullet}$, $\bar{B}_{\bullet j}$ and $\bar{B}_{\bullet\bullet}$ the row, column and grand means, which enforces both marginal constraints simultaneously and renders the rewiring channel orthogonal to the abundance channels. This orthogonality is what allows $\beta^{\pi}$ to isolate genuine preference shifts rather than double-counting species individual effects-driven changes already absorbed by $\beta^{p}$ and $\beta^{a}$.

To avoid confounding the urbanisation contrast with species turnover, the contrasts are estimated only for species present in *both* groups. Writing $\mathcal{C}_p$ and $\mathcal{C}_a$ for the common plant and pollinator sets, the parameters $\beta_i^{p}$, $\beta_j^{a}$ and $\beta_{ij}^{\pi}$ are defined on $\mathcal{C}_p$, $\mathcal{C}_a$ and $\mathcal{C}_p \times \mathcal{C}_a$, and the sum-to-zero and double-centring constraints are applied over these sets; species seen in a single group keep their baseline effect but receive no low$\rightarrow$high contrast.

All effects use the same non-centred parametrization as the rest of the model to improve the sampling geometry. The overall shift is weakly regularised, $\beta_{\gamma} \sim \mathcal{N}(0, 1)$, and each contrast channel is given a hierarchical scale shrunk toward zero,

$$\beta_i^{p} \sim \mathcal{N}(0, \sigma_{\beta^p}^2), \quad \beta_j^{a} \sim \mathcal{N}(0, \sigma_{\beta^a}^2), \quad \beta_{ij}^{\pi} \sim \mathcal{N}(0, \sigma_{\beta^{\pi}}^2), \tag{S13}$$

with half-normal hyperpriors $\sigma_{\beta^p}, \sigma_{\beta^a}, \sigma_{\beta^\pi} \sim \mathcal{N}^+(0, 0.5^2)$. The sum-to-zero raw effects are additionally rescaled by $\sqrt{n/(n-1)}$ so that each element retains unit marginal prior variance before multiplication by its scale, keeping the per-element prior scale equal to $\sigma_{\beta.}$ despite the constraint. These half-normal priors concentrate mass near zero, encoding a default of "no urbanisation effect" and letting the data pull each channel away from it only when evident, which provides principled regularisation in the small-sample two-group design.

From the fitted contrasts we recover group-specific summaries on the natural scale. Species activities are reported as $\exp(\bar{p}_i)$ and $\exp(\bar{p}_i + \beta_i^p)$ for the low and high groups (analogously for pollinators), and preference networks as $\exp(\pi_{ij})$ and $\exp(\pi_{ij} + \beta_{ij}^\pi)$. The channel-specific scale parameters $\sigma_{\beta^p}, \sigma_{\beta^a}, \sigma_{\beta^\pi}$ quantify the magnitude of the urbanisation response along each ecological axis, indicating whether urbanisation acts primarily through shifts in species activity or through network rewiring.

### B. Posterior summaries

The variance parameters not tied to the covariate summarise the community as observed across sites and months (Table S6). Residual site variation is modest ($\Sigma_\gamma = 0.19$, 95% PCI $[0.08, 0.40]$), indicating that the low/high contrast captures most of the between-site structure. Species baseline activity is small and only weakly identified ($\sigma_p = 0.21$ $[0.01, 0.56]$; $\sigma_a = 0.30$ $[0.02, 0.51]$), with most of it absorbed by the time-varying terms ($\Sigma_{p,t} = 0.74$ $[0.57, 0.94]$; $\Sigma_{a,t} = 0.35$ $[0.20, 0.51]$); temporal fluctuation, not stable species identity, dominates activity. The two time-varying terms differ by roughly a factor of two, so plant activity varies about twice as much across visits as pollinator activity, consistent with sampling over the flowering season. Preference heterogeneity is by far the largest variance component and is tightly estimated ($\sigma_\pi = 1.37$ $[1.24, 1.52]$), confirming the strong, well-resolved preference structure highlighted in the main text. Activity is moderately persistent between consecutive visits ($\rho = 0.53$ $[0.26, 0.73]$), well away from both 0 and 1, which justifies keeping visits separate rather than pooling them.

| Parameter | Mean | SD | 2.5% | 50% | 97.5% |
|---|---|---|---|---|---|
| $\Sigma_\gamma$ | 0.192 | 0.082 | 0.080 | 0.177 | 0.399 |
| $\sigma_p$ | 0.212 | 0.156 | 0.008 | 0.185 | 0.563 |
| $\sigma_a$ | 0.295 | 0.123 | 0.024 | 0.308 | 0.512 |
| $\sigma_\pi$ | 1.372 | 0.072 | 1.240 | 1.371 | 1.521 |
| $\Sigma_{p,t}$ | 0.743 | 0.094 | 0.565 | 0.740 | 0.938 |
| $\Sigma_{a,t}$ | 0.345 | 0.078 | 0.199 | 0.341 | 0.507 |

TABLE S6. Posterior summary statistics of global variance parameters from the multi-observation model with urbanisation covariate fitted to the dataset of [4].

### C. Convergence diagnostics

For the urbanisation case study, all convergence diagnostics were satisfactory (Table S7). Due to the presence of the autoregressive temporal component, we ran 1500 post-warmup iterations per chain (4 chains). The rank-normalized split-$\widehat{R}$ did not exceed 1.01 for any parameter, and both bulk- and tail-effective sample sizes remained above the target of 400 throughout (smallest bulk-ESS 512, for the temporal-correlation parameter $\rho$). In particular, the urbanisation channels, the differential-abundance effects $\beta_i^p, \beta_j^a$ and the rewiring effects $\beta_{ij}^\pi$, were well sampled (bulk-ESS $\geq$ 3300), as were the pairwise preferences $\pi_{ij}$ (bulk-ESS $\geq$ 2200). Stan's diagnostic checks reported no divergent transitions, no saturation of the maximum tree depth, and satisfactory energy diagnostics (E-BFMI).

### D. Network metrics

To assess network structure difference of plant-pollinator communities in the two urbanisation levels, we consider weighted connectance [1] and interaction diversity [8].

*Weighted Connectance.* Weighted connectance is a measure of how densely and how evenly the matrix is filled, accounting for cell values. Specifically, let $M \in \mathbb{R}_{\geq 0}^{n_p \times n_a}$ be the *interaction* matrix between $n_p$ plant species and $n_a$ pollinator species, with $m_{ij}$ denoting either the observed visit count between plant $i$ and pollinator $j$, or the inferred preference between the pair. Let $m_\bullet = \sum_{ij} m_{ij}$ the total number of visits. For each row $i$, we define the normalised visit profile $p_{ij} = m_{ij}/m_{i\bullet}$ and its Shannon

| Parameter | $\widehat{R}$ | bulk-ESS | tail-ESS |
|---|---|---|---|
| $\bar{\gamma}$ | 1.000 | 2813 | 3903 |
| $u_s$ | 1.000 | 4201–4716 | 4077–4869 |
| $\bar{p}_i$ | 1.000 | 1380–9737 | 1581–4349 |
| $\bar{a}_j$ | 1.000 | 851–11930 | 1584–5069 |
| $\eta^p_{i,t}$ | 1.000 | 1326–13624 | 1866–5572 |
| $\eta^a_{j,t}$ | 1.000 | 945–14625 | 1504–5197 |
| $\rho$ | 1.000 | 512 | 1313 |
| $\pi_{ij}$ | 1.000–1.010 | 2267–17750 | 3042–5272 |
| $\beta^\gamma$ | 1.000 | 2663 | 3377 |
| $\beta^p_i$ | 1.000 | 5147–10653 | 3945–5068 |
| $\beta^a_j$ | 1.000 | 3373–13663 | 3275–4931 |
| $\beta^\pi_{ij}$ | 1.000–1.010 | 5334–16598 | 3363–5201 |
| $\sigma_{\text{site}}$ | 1.000 | 2993 | 4201 |
| $\sigma_p$ | 1.010 | 986 | 1778 |
| $\sigma_a$ | 1.000 | 737 | 1109 |
| $\sigma_\pi$ | 1.000 | 2412 | 4284 |
| $\Sigma_{p,t}$ | 1.000 | 1087 | 2423 |
| $\Sigma_{a,t}$ | 1.000 | 886 | 2045 |
| $\sigma_{\beta^p}$ | 1.000 | 2361 | 2294 |
| $\sigma_{\beta^a}$ | 1.000 | 1743 | 1910 |
| $\sigma_{\beta^\pi}$ | 1.000 | 1266 | 1453 |

TABLE S7. Convergence diagnostics for the urbanisation case study, reported by parameter block. Values are ranges (minimum–maximum) across the parameters in each block, except for scalar parameters. The blocks comprise the baseline intensity $\bar{\gamma}$ and site effects $u_s$; the core species effects $(\bar{p}_i, \bar{a}_j)$ with their temporal deviations $(\eta^p_{i,t}, \eta^a_{j,t})$; the temporal-correlation parameter $\rho$; the pairwise preferences $\pi_{ij}$; the urbanisation channels ($\beta^\gamma$, $\beta^p_i$, $\beta^a_j$, and $\beta^\pi_{ij}$); and the random-effect scales ($\sigma_{\text{site}}, \sigma_p, \sigma_a, \sigma_\pi$, the temporal-deviation scales $\Sigma_{p,t}$, $\Sigma_{a,t}$, and the urbanisation-channel scales $\sigma_{\beta^p}, \sigma_{\beta^a}, \sigma_{\beta^\pi}$). All $\widehat{R} \leq 1.01$ and all bulk and tail effective sample sizes exceed 400.

entropy $H_i = -\sum_j p_{ij} \log_2 p_{ij}$; analogously for each column $j$. The effective number of partners of species $i$ is then $2^{H_i}$, and the *linkage density* is the interaction-weighted average across both guilds:

$$\text{LD} = \frac{1}{2}\left(\sum_i \frac{m_{i\bullet}}{m_\bullet} 2^{H_i} + \sum_j \frac{m_{\bullet j}}{m_\bullet} 2^{H_j}\right). \tag{S14}$$

Weighted connectance is then $C_w = \text{LD}/(n_p + n_a)$. A network where every species interacts with every other species with equal *weight* yields $C_w = 0.5$, while a network dominated by a few strong interactions between a few species yields a low $C_w$. It differs from binary connectance, which simply counts the proportion of non-zero entries $|\{(i,j) : m_{ij} > 0\}|/(n_p \cdot n_a)$, in that two networks with identical link presence patterns but different visit or preference distributions will have the same binary connectance but different weighted connectance.

*Interaction Diversity.* Interaction diversity summarises an interaction matrix by the effective number of interactions it contains. We define the probability associated with the pair $(i, j)$ as $p_{ij} = m_{ij}/m_\bullet$, so that the entries of $M$ are treated as a distribution of interaction weight over pairs. The Shannon entropy of this distribution is

$$H = -\sum_{i,j:\, m_{ij}>0} p_{ij} \log p_{ij}, \tag{S15}$$

and interaction diversity is its exponential, $D = e^H$, the Hill number of order $q = 1$. Conceptually, $D$ answers the question: how many pairs, each carrying equal weight, would reproduce the entropy of the observed distribution? A network of $k$ pairs of equal weight gives $D = k$, whereas concentrating all the weight on a single pair gives $D = 1$ regardless of how many other pairs are present. Interaction diversity thus captures in one number both the richness of interactions (how many pairs interact at all) and how evenly weight is spread across them, with a natural effective-number interpretation.

#### E. Interaction $\beta$-diversity decomposition

We further characterise the low–high urbanisation contrast by decomposing interaction $\beta$-diversity with the framework introduced in [10]. Given the networks corresponding to the two classes, whether these comes from observed counts or inferred

| Metric | Observed counts | Latent preferences |
|---|---|---|
| $\beta_{\mathrm{WN}}$ | 0.58 | 0.66 [0.62, 0.70] |
| $\beta_{\mathrm{OS}}$ (rewiring) | 0.50 | 0.20 [0.10, 0.29] |
| $\beta_{\mathrm{ST}}$ (turnover) | 0.08 | 0.46 [0.40, 0.54] |
| Rewiring fraction | 0.87 | 0.30 [0.16, 0.42] |

TABLE S8. Interaction $\beta$-diversity decomposition [10] computed on the inferred latent preferences versus the observed counts. Total dissimilarity $\beta_{WN}$ splits into a rewiring component among shared species ($\beta_{OS}$) and a species-turnover component ($\beta_{ST}$), with the rewiring fraction $\beta_{OS}/\beta_{WN}$. Values related to preferences are shown as posterior medians with 95% posterior credible interval. Rewiring accounts for the majority of whole-network dissimilarity on the observed counts (0.87) but drops to 0.30 once abundance is controlled for by measuring on the latent preferences.

preferences, the decomposition rests on the additive identity

$$\beta_{WN} = \beta_{ST} + \beta_{OS}, \tag{S16}$$

where $\beta_{WN}$ is a dissimilarity measured over all interactions, $\beta_{OS}$ over the sub-matrix of species present in both networks (shared plants crossed with shared pollinators), and $\beta_{ST}$ is obtained by subtraction, $\beta_{ST} = \beta_{WN} - \beta_{OS}$. Species presence is computed from non-zero marginal totals of the matrix considered. Because $\beta_{OS}$ looks only at species common to both networks, it captures the extent to which shared partners are wired differently (rewiring), while $\beta_{ST}$ represents the dissimilarity traceable to species that occur in one network only (turnover). We summarise the balance of the two components with the rewiring fraction $\beta_{OS}/\beta_{WN}$.

Since our networks are weighted, we following [10] and measure each term with the quantitative (Bray–Curtis) index. For two weight vectors $x$ and $y$, this is given by:

$$\mathrm{BC}(x, y) = \frac{\sum_i |x_i - y_i|}{\sum_i (x_i + y_i)}. \tag{S17}$$

Ranging from 0 (identical weight profiles) to 1 (no interactions in common), this expresses the summed absolute weight difference as a fraction of the total weight, so that mismatches on heavily weighted links count proportionally more. Each matrix is normalised to sum to one before BC is evaluated, and the shared sub-block is renormalised to sum to one before $\beta_{OS}$ is computed, such that dissimilarity reflects how weights (visits or preferences) are distributed rather than the totals.

Values computed for observed counts and latent preferences are reported in Table S8.

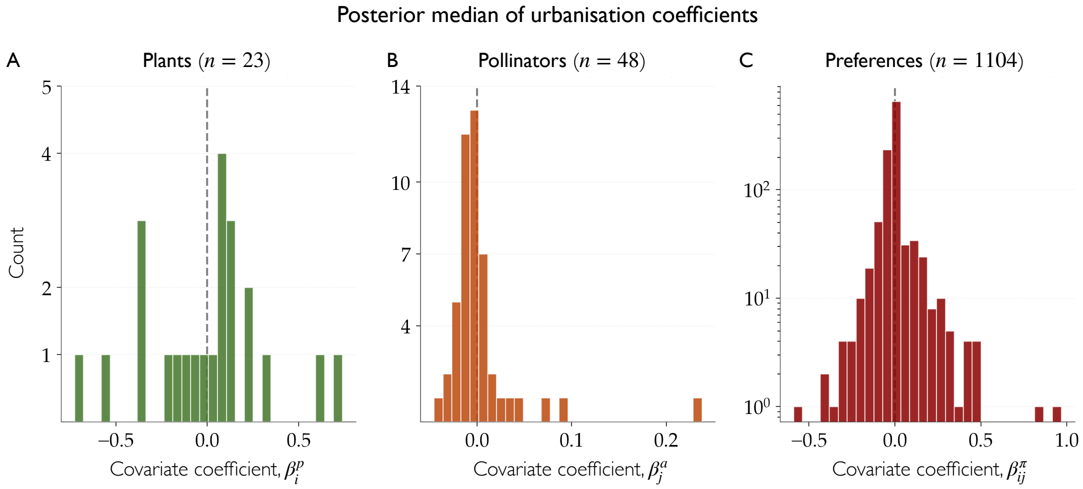


FIG. S13. **Distribution of posterior median of urbanisation coefficients.** Histograms of the posterior median of urbanisation coefficient ($x$-axis) for each channel of the model: plant species-level effects ($\beta_i^p$, **A**), pollinator species-level effects ($\beta_j^a$, **B**), and pairwise interaction preferences ($\beta_{ij}^\pi$, **C**). Coefficients are defined along the urbanisation gradient (low $\rightarrow$ high), so that positive and negative values respectively indicate quantities that increase and decrease with urbanisation; the dashed grey line marks zero (no effect). The $y$-axis gives the number of species (**A**, **B**) or species pairs (**C**) falling in each bin; note the logarithmic $y$-axis in panel **C**. Coefficients are shown only for species recorded at both low- and high-urbanisation sites, giving $n = 23$ plants, $n = 48$ pollinators, and $n = 1104$ preference pairs.

[1] L.-F. Bersier, C. Banašek-Richter, and M.-F. Cattin. Quantitative descriptors of food-web matrices. *Ecology*, 83(9):2394–2407, 2002.

[2] G. E. Box. Sampling and bayes' inference in scientific modelling and robustness. *Journal of the Royal Statistical Society Series A: Statistics in Society*, 143(4):383–404, 1980.

[3] S. R. Cook, A. Gelman, and D. B. Rubin. Validation of software for bayesian models using posterior quantiles. *Journal of Computational and Graphical Statistics*, 15(3):675–692, 2006.

[4] A. Fisogni, N. Hautekèete, Y. Piquot, M. Brun, C. Vanappelghem, M. Ohlmann, M. Franchomme, C. Hinnewinkel, and F. Massol. Seasonal trajectories of plant-pollinator interaction networks differ following phenological mismatches along an urbanization gradient. *Landscape and Urban Planning*, 226:104512, 2022.

[5] A. Gelman, A. Vehtari, D. Simpson, C. C. Margossian, B. Carpenter, Y. Yao, L. Kennedy, J. Gabry, P.-C. Bürkner, and M. Modrák. Bayesian workflow. *arXiv preprint arXiv:2011.01808*, 2020.

[6] C. J. Geyer. Introduction to markov chain monte carlo. In S. Brooks, A. Gelman, G. L. Jones, and X.-L. Meng, editors, *Handbook of Markov Chain Monte Carlo*, pages 3–48. Chapman & Hall/CRC, 2011.

[7] M. D. Hoffman and A. Gelman. The no-u-turn sampler: Adaptively setting path lengths in hamiltonian monte carlo. *Journal of Machine Learning Research*, 15(47):1593–1623, 2014.

[8] L. Jost. Entropy and diversity. *Oikos*, 113(2):363–375, 2006.

[9] D. N. Moriasi, J. G. Arnold, M. W. Van Liew, R. L. Bingner, R. D. Harmel, and T. L. Veith. Model evaluation guidelines for systematic quantification of accuracy in watershed simulations. *Transactions of the ASABE*, 50(3):885–900, 2007.

[10] T. Poisot, E. Canard, D. Mouillot, N. Mouquet, and D. Gravel. The dissimilarity of species interaction networks. *Ecology letters*, 15(12): 1353–1361, 2012.

[11] E. L. Rezende, J. E. Lavabre, P. R. Guimarães, P. Jordano, and J. Bascompte. Non-random coextinctions in phylogenetically structured mutualistic networks. *Nature*, 448, 2007.

[12] T. Säilynoja, P.-C. Bürkner, and A. Vehtari. Graphical test for discrete uniformity and its applications in goodness of fit evaluation and multiple sample comparison. *arXiv preprint arXiv:2103.10522*, 2021.

[13] S. Talts, M. Betancourt, D. Simpson, A. Vehtari, and A. Gelman. Validating bayesian inference algorithms with simulation-based calibration. *arXiv preprint arXiv:1804.06788*, 2018.

[14] A. Vehtari, A. Gelman, D. Simpson, B. Carpenter, and P.-C. Bürkner. Rank-normalization, folding, and localization: An improved $\widehat{R}$ for assessing convergence of MCMC. *Bayesian analysis*, 16(2):667–718, 2021.
